# Stealth replication of SARS-CoV-2 Omicron in the nasal epithelium at physiological temperature

**DOI:** 10.1101/2025.05.03.652024

**Authors:** Barbara F. Fonseca, Rémy Robinot, Vincent Michel, Akram Mendez, Samuel Lebourgeois, Chloé Chivé, Raphaël Jeger-Madiot, Roshan Vaid, Vincent Bondet, Elizabeth Maloney, Florence Guivel-Benhassine, Olivier Schwartz, Darragh Duffy, Tanmoy Mondal, Samy Gobaa, Lisa A. Chakrabarti

## Abstract

The COVID-19 pandemic was marked by successive waves of SARS-CoV-2 variants with distinct properties. The Omicron variant that emerged in late 2021 showed a major antigenic shift and rapidly spread worldwide. Since then, Omicron-derived variants have maintained their global dominance, for reasons that remain incompletely understood. We report that the original Omicron variant BA.1 evolved several traits that converged in facilitating viral spread. First, Omicron displayed an early replicative advantage over previous variants when grown in a reconstructed nasal epithelium model based on primary human cells. The increase in Omicron replication was more marked at the 33°C temperature characteristic of human nasal passages, resulting in a physiologically relevant advantage. Omicron also caused a decrease in epithelial integrity, as measured by transepithelial electrical resistance and caspase-3 activation. Furthermore, Omicron caused a more marked loss of motile cilia at 33°C than other variants, suggesting a capacity to impair mucociliary clearance. RNAseq analysis showed that Omicron induced a broad transcriptional downregulation of ciliary genes but only a limited upregulation of host innate defense genes at 33°C. The lower production of type I and type III interferons in epithelia infected by Omicron compared to those infected by the Delta variant, at 33°C as well as 37°C, confirmed the increased capacity of Omicron to evade the innate antiviral response. Thus, Omicron combined replication speed, motile cilia impairment, and limited induction of innate antiviral responses when propagated in reconstructed nasal epithelia at physiological temperature. Omicron has the capacity to propagate efficiently but stealthily in the upper respiratory tract, which likely contributed to the evolutionary success of this SARS-CoV-2 variant.

**AUTHOR SUMMARY:** The COVID-19 pandemic was initially characterized by a rapid succession of viral variants that emerged independently of each other, with each of these variants outcompeting previous ones and rising to regional or global dominance. A major evolutionary shift occurred in late 2021, with the emergence of the highly divergent Omicron BA.1 variant. Since then, all the dominant SARS-CoV-2 variants have been derived from Omicron, for reasons that remain incompletely understood. In this study, we chose to compare the replication of SARS-CoV-2 variants in a human nasal epithelium model grown at 37°C but also at 33°C, a more physiological temperature that approximates that found in the nasal cavity. In this model, Omicron showed an early replicative advantage that was more marked at nasal physiological temperature. Omicron also markedly impaired the layer of motile cilia that normally contributes to the clearance of inhaled particles from the nasal mucosa. Even though it caused tissue damage, Omicron triggered only a minimal antiviral interferon response from epithelia grown at 33°C. Thus, Omicron has the capacity to propagate rapidly but stealthily in the nasal epithelium at physiological temperature, which helps account for the efficient dissemination of this variant worldwide.

## INTRODUCTION

The COVID-19 pandemic was characterized by sequential evolutionary shifts, with a rapid succession of viral variants rising to global dominance [1][2]. The initial SARS-CoV-2 Wuhan strain that emerged in late 2019 was replaced in the first semester of 2020 by a variant with higher infectivity, due to the presence of a D614G mutation in the S1 surface subunit of the Spike protein [3]. Starting from late 2020, a series of variants termed Alpha, Beta, Gamma, Delta, and Omicron emerged successively and independently of each other, with each of these variants of concern (VOCs) outcompeting previous variants. The Delta variant, which reached worldwide dominance in spring 2021, was associated with increased disease severity, which was in part ascribed to a higher fusogenic capacity of its Spike glycoprotein [4][5][6]. A major evolutionary shift occurred in late 2021, with the emergence of the Omicron BA.1 variant that differed by more than 32 amino acids in the Spike glycoproteins from previous strains [7]. Since then, all the dominant SARS-CoV-2 variants have been phylogenetically derived from Omicron, with adaptation to preexisting immunity becoming a major selective pressure for variant replacement [8][9].

Why Omicron-derived variants have maintained their global dominance remains incompletely understood. Epidemiological studies have documented a marked increase in viral fitness upon Omicron emergence, as measured by rates of variant spread over time [10]. While escape from preexisting neutralizing antibodies are thought to be a key factor to explain the Omicron surge, there are signs that this variant is also intrinsically efficient at transmission from human to human [1]. In particular, longitudinal analyses of nasopharyngeal swabs showed that detectable viral replication in the nasal mucosa occurs earlier for the Omicron than the Delta variant [11]. Consistent with this observation, Omicron shows faster replication kinetics than previous variants in human airway explants [12] and in reconstructed epithelial cultures derived from primary human nasal cells [13][14][15].

While the Omicron variant spread efficiently, it proved slightly less pathogenic than the preceding Delta variant, as measured by hospitalization rates and pneumonia severity in individuals who were not previously vaccinated or infected by SARS-CoV-2 [16][17]. These epidemiological observations are in agreement with findings in animal models, with a lower degree of lung dysfunction induced by Omicron in infected golden Syrian hamsters, ferrets, and non-human primates [18][19][13][20]. Of note, while most animal models show an attenuated Omicron phenotype, they often do not reproduce the rapid viral replication observed in the human nasal mucosa. It is also relevant that while Omicron replicates faster than Delta in human bronchus explants, it replicates more slowly than Delta in human lung explants [12] and alveolar organoids [21][22]. This differential tropism fits with the notion of a bias towards the upper over the lower respiratory tract, possibly accounting for a lower pathogenic potential of Omicron.

Different mechanisms have been proposed to account for the increased replication capacity of Omicron in human airway epithelia, including a higher affinity of Spike for the ACE-2 receptor [23][24], or a better interaction of Spike with the negatively charged glycocalyx at the cellular surface [25], leading to an overall better attachment of viral particles to primary ciliated epithelial cells [26]. Several studies have also documented that Omicron entry pathway differs from that of previous variants, with a lower reliance on the surface serine protease TMPRSS2 to release the fusion peptide and an expanded usage of other proteases [27][28][14]. In addition, an improved capacity to evade the innate antiviral response has also been reported for Omicron, which could contribute to its rapid replication kinetics [29][30][31]. To further address these questions, we systematically compared the replication of Omicron and previous variants in a reconstructed human nasal epithelium model grown at the air-liquid interface (ALI). This model recapitulates key features of the nasal mucosa, including a pseudostratified epithelium structure, the presence of diverse epithelial cell types (ciliated, goblet, and basal cells), and the competence for mucociliary clearance function [32]. In addition to its barrier function, the nasal epithelium has also an air conditioning function, warming and moistening inspired air to maintain the internal milieu of the lungs [33]. This results in a temperature gradient from the nares to the nasopharynx, with 31-34°C typically measured at the posterior end of the nasal cavity in humans [34]. Therefore, we chose to compare SARS-CoV-2 variant replication at 37°C but also at 33°C, a more physiological temperature that approximates that found in the nasal cavity. We report that the Omicron replicative advantage is more marked at the temperature found in nasal passages, as this variant manages to spread efficiently and induce motile cilia damage while triggering only a minimal innate interferon (IFN) response from primary epithelial cells at 33°C. These findings help account for the rapid and efficient dissemination of the Omicron variant worldwide.

## RESULTS

### The replicative advantage of Omicron in nasal epithelial cells is more marked at a physiologically relevant temperature

For infection experiments, we used reconstructed human nasal epithelia generated from pooled human donors (MucilAir^TM^, Epithelix), to minimize donor-dependent variability. Prior to infection, cultures maintained at the air-liquid interface (ALI) were checked for motile cilia activity and mucus production, ensuring the proper differentiation of a pseudostratified epithelium. Infectious inocula with the different SARS-CoV-2 variants were normalized based on viral RNA content and infections were monitored for 4 days.

In nasal epithelial cultures grown at 37°C, all the viruses tested showed an efficient viral replication, including the ancestral Wuhan strain, the early variant D614G, and the later variants Alpha, Delta, and Omicron BA.1 (Fig. 1A), as well as the Beta and Gamma variants (Supp. Fig. S1A, C). Viral replication reached a plateau at day 2 (D2), except for the Omicron BA.1 variant, which reached close to plateau values at D1. Interestingly, the replicative advantage of Omicron BA.1 became more apparent when the epithelial cultures were grown at 33°C, a temperature more relevant to nasal physiology (Fig. 1B and Supp. Fig. S1B, D). Comparison of viral RNA copy number values at D1 showed a trend for higher replication of BA.1 compared to the D614G reference variant at 37°C (not significant; Fig. 1C), and confirmed that BA.1 replication was significantly higher than that of D614G at 33°C (Fig. 1D, P=0.016). The replicative advantage of Omicron was apparent at early time points, but the replication of other VOCs caught up with that of BA.1 at the later D4 time point, possibly due to a limitation in target cell availability.

**Figure 1:**
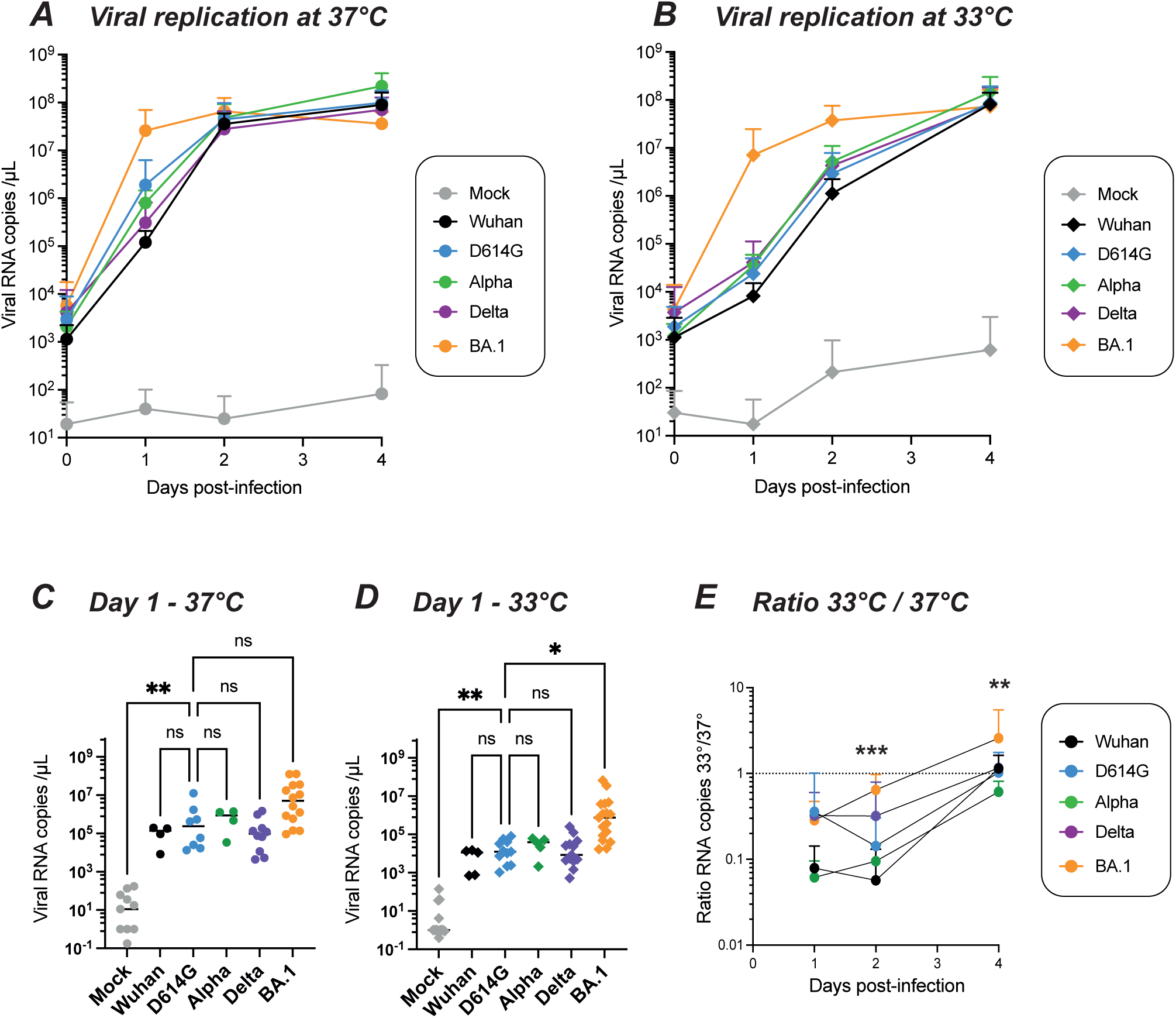
Temperature-dependent replication of SARS-CoV-2 variants in reconstructed human nasal epithelia. The viral load in apical supernatants was quantified by RT-qPCR for samples infected by the following SARS-CoV-2 lineages: Wuhan, D614G, Alpha, Delta and Omicron BA.1. (A, B) Kinetics of viral replication at 37°C (A) and at 33°C (B), with means and SD reported (n=4 to 14 independent samples per point). (C,D) Comparison of apical viral loads measured at 1 day post infection (dpi) at 37°C (C) and 33°C (D). Medians are reported. (E) The ratio of the number of viral RNA copies measured at 33°C to that measured at 37°C is shown at different dpi, with means and SD reported (n=4 to 14 independent samples per point).(C, D, E) Statistical comparisons were done between the D614G reference and all other variants or the mock condition, using the Kruskal-Wallis test with Dunn’s correction; ns: not significant; * P<0.05; ** P<0.01; *** P<0.001. In (E), the only significant differences in ratios were seen between BA.1 and D614G at 2 dpi and 4 dpi.

Computation of the ratio of viral RNA copies at 33°C over those at 37°C showed that viral replication tended to be slower at the 33°C temperature for all the viruses tested, with ratios below 1 in most cases, except for Omicron BA.1 at D4 (Fig. 1E, Supp. Fig. 1E). In addition, Omicron BA.1 was the only variant which showed a ratio significantly higher than that of the D614G reference strain at D2 (P<0.001) and D4 (P<0.01), emphasizing the efficient replication of the Omicron variant at a temperature more physiologically relevant for nasal infection.

### Omicron infection perturbs the epithelial barrier at nasal physiological temperature

The trans-epithelial electrical resistance (TEER) measured between the upper and lower compartment of transwell cultures was used to evaluate the integrity of the epithelial barrier. All the variants tested induced a decrease in TEER at D4 as compared to mock-infected epithelia (Fig. 2A-B), indicating that SARS-CoV-2 perturbs epithelial barrier function. The extent of TEER decrease was comparable for all variants tested at 37°C, with a median decrease between 1.5 and 1.9-fold (Fig. 2A). In contrast, at the lower temperature of 33°C, the Omicron BA.1 variant caused a more marked TEER decrease than the reference D614G variant (median decrease of 2.5x vs 1.3x; P=0.003; Fig. 2B). Thus, the Omicron variant proved particularly damaging for nasal epithelial integrity at the physiologically relevant temperature of 33°C.

**Figure 2:**
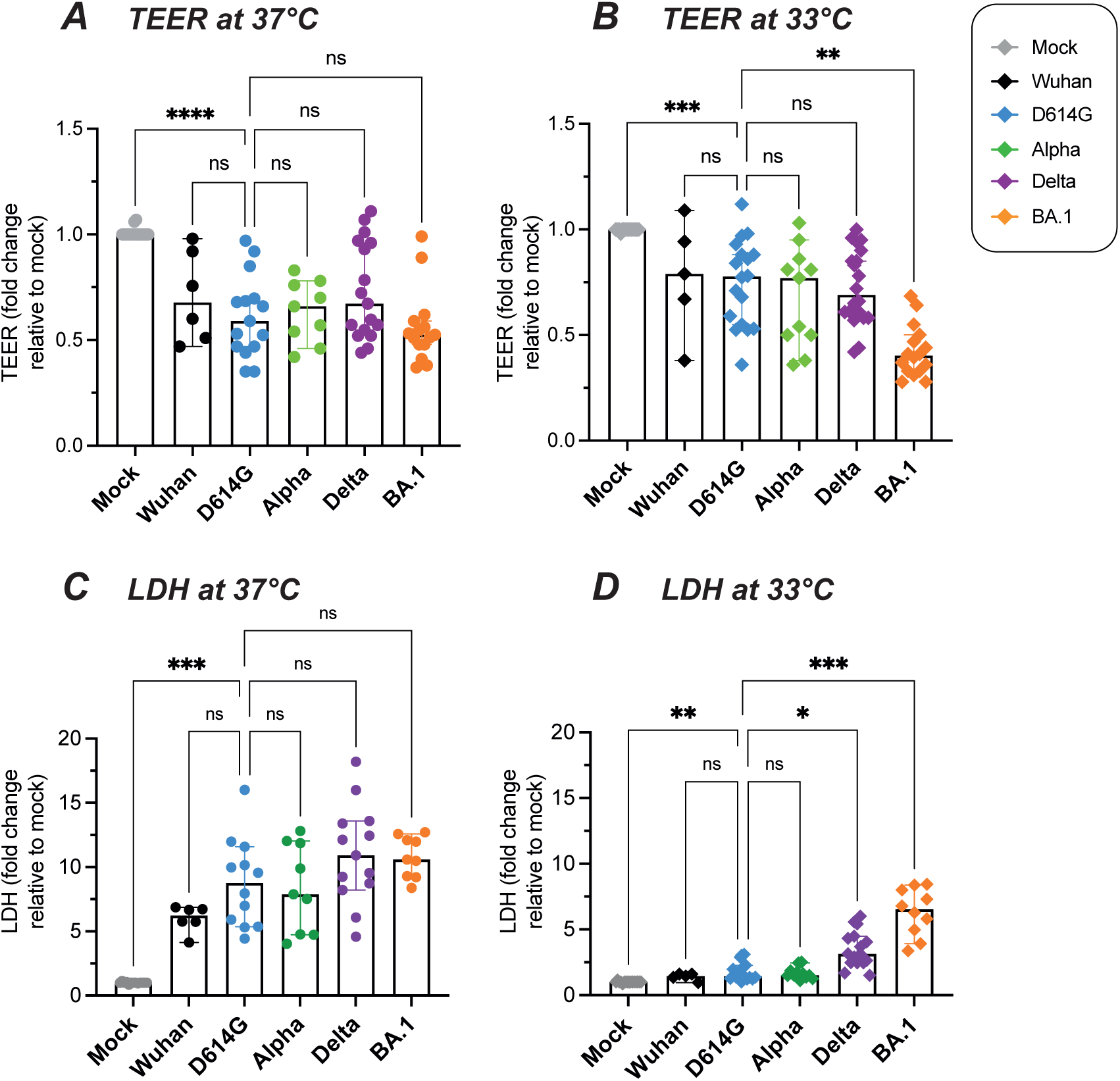
SARS-CoV-2 variants impair epithelial integrity. (A, B) Epithelial barrier function was assessed at 37°C (A) and 33°C (B) by measuring the trans-epithelial electric resistance (TEER) at 4 dpi. Relative TEER values reported to the Mock condition are shown. (C, D) The cytopathic effect was assessed at 37°C (C) and 33°C (D) by measuring the release of lactate dehydrogenase (LDH) in apical supernatants at 4 dpi. Relative LDH concentrations reported to the Mock condition are shown. (A-D) Medians and 95% CI are reported. Statistical comparisons were done between the D614G reference and all other variants or the Mock condition, using the Kruskal-Wallis test with Dunn’s correction; ns: not significant; * P<0.05; ** P<0.01; *** P<0.001; **** P<0.0001.

We next asked whether epithelial damage was associated with cell death, by measuring the apical release of lactate dehydrogenase (LDH) at D4 (Fig. 2 C-D). All the variants tested induced an increase in LDH release, confirming the cytopathic effect of SARS-CoV-2 infection. The extent of LDH release did not differ significantly between the variants tested at 37°C (Fig. 2C). LDH release was overall lower at 33°C (Fig. 2D). However, at this lower temperature, the Delta and Omicron BA.1 variants showed a significantly increased cytopathic effect compared to the reference D614G variant (P=0.004 for Delta; P=0.0006 for BA.1). It was noteworthy that the Delta variant showed a clear cytopathic effect in the absence of a replicative advantage in epithelial cultures, a phenomenon that may be ascribed to the high fusogenicity of this variant [5][35].

To further investigate the cytopathic effect induced by the different SARS-CoV-2 variants, we evaluated by immunofluorescence the expression of the cleaved active form of caspase-3, an apoptosis effector protein. Cleaved caspase-3 was induced by infection at D4, and more so at 37°C than at 33°C (Fig. 3A). Fluorescence quantification showed that Omicron BA.1 was the most efficient at inducing cleaved caspase-3 at both temperatures (P<0.05 in both cases), while Delta showed a non-significant trend for increase as compared to the D614G and Wuhan strains (Fig. 3 B-C). The Spike and cleaved caspase-3 signals colocalized in some but not all infected cells, pointing to caspase-3 activation in infected cells but also possibly in bystander cells. It was interesting to note that large Spike+ cells corresponding to syncytia appeared more prominent for the Delta variant (Fig. 3A, 3rd row), consistent with the highly fusogenic nature of this variant. Taken together, these findings documented a gradation in the caspase-3-mediated cytopathic effect, with a maximal induction by Omicron BA.1, an intermediate induction by Delta, and a lower induction by other variants. Of note, Omicron significantly perturbed epithelial integrity and viability even at the physiological temperature of 33°C.

**Figure 3:**
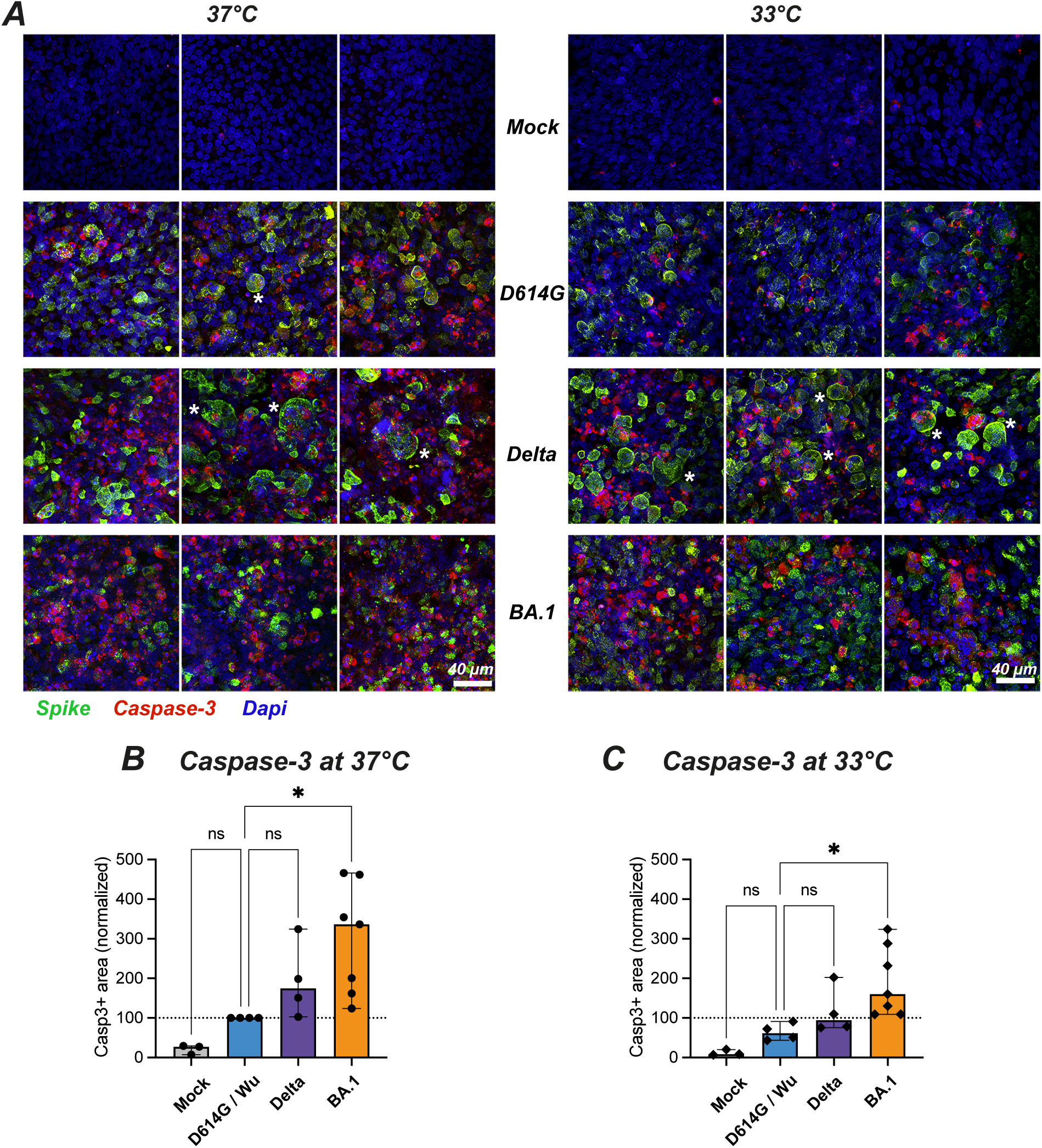
Temperature-dependent cytopathic effects induced by different SARS-CoV-2 variants. (A) Reconstructed nasal epithelia were mock-infected (top line) or infected with different SARS-CoV-2 variants (D614G, Delta, Omicron BA.1) at 37°C (left) or 33°C (right) were fixed at 4 dpi and immunolabeled for cleaved caspase-3 (red), Spike (green) and nuclei (Dapi, blue). The scale bar represents 40 µm. The * symbol labels large syncytia. (B-C) Quantification of cleaved caspase 3 positive area (casp3+) at 37°C (B) and 33°C (C), with values normalized to the D614G or Wuhan values (D614G/Wu) at 37°C. Medians and 95% CI are reported. Statistical comparisons were done between the D614G/Wu reference and all other variants or the Mock condition, using the Kruskal-Wallis test with Dunn’s correction; ns: not significant; * P<0.05.

### Omicron infection induces the loss of motile cilia at nasal physiological temperature

We next evaluated the impact of the different viral variants on the motile cilia layer that covers airway epithelia. Reconstructed nasal epithelia collected at D4 post-infection were labeled for the acetylated form of α−tubulin, a tubulin isoform that is enriched in the axoneme of motile cilia. Immunofluorescence analysis showed that the three variants tested induced a loss of motile cilia at 37°C (Fig. 4A, left), with a decrease that was significant only for Omicron BA.1 (Fig. 4B, P=0.026). The loss of cilia proved less marked at 33°C (Fig. 4A, right), with again a significant decrease seen only for BA.1 (Fig. 4C; P=0.002).

**Figure 4:**
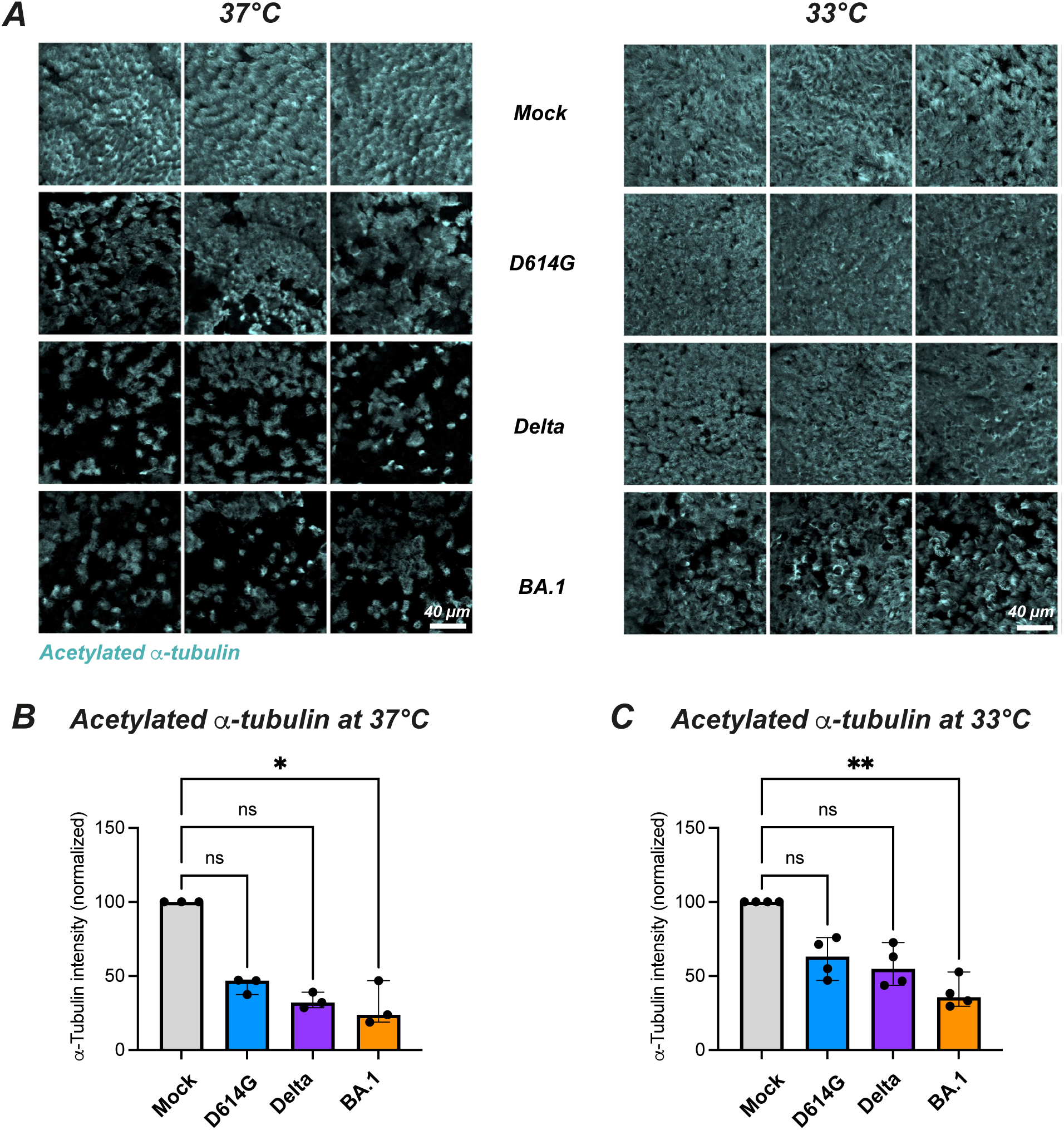
SARS-CoV-2 variants perturb the motile cilia layer in a temperature-dependent fashion. (A) Reconstructed nasal epithelia were mock-infected (top line) or infected with different SARS-CoV-2 variants (D614G, Delta, Omicron BA.1) at 37°C (left) or 33°C (right) were fixed at 4 dpi and immunolabeled for acetylated α-tubulin (cyan), which preferentially detects motile cilia. The scale bar represents 40 µm. (B-C) Quantification of the area positive for acetylated α-tubulin at 37°C (B) and 33°C (C), with values normalized to the mock-infected condition. Medians and 95% CI are reported. Statistical comparisons were done between the mock-infected sample and all the SARS-CoV-2 variant infected samples, using the Kruskal-Wallis test with Dunn’s correction; ns: not significant; * P<0.05; ** P<0.01.

Examination of the infected epithelia by scanning electron microscopy confirmed the disruption of the motile cilia layer by infection at D4, with the presence of multiple rounded cells with partial or complete deciliation, especially at 37°C (Sup. Fig. S2). The observation of membrane protrusions surrounding cilia suggested that engulfment may contribute to cilia loss. Many of the rounded deciliated cells appeared pushed to the surface of the epithelial layer, consistent with the extrusion of dying epithelial cells. Rounded imprints corresponding to missing cells were prominent in BA.1 infected epithelia. The detection of blebbing cells also pointed to the occurrence of apoptosis. Overall, infection by D614G, Delta and BA.1 all perturbed the ciliary layer at 37°C, while changes were more prominent for the BA.1 variant at 33°C.

### Omicron induces a broad transcriptional downregulation of ciliary genes but only a limited upregulation of host defense genes at nasal physiological temperature

To identify the transcriptional changes that may contribute to epithelial perturbation, we performed bulk RNAseq analysis on epithelia at D2 post-infection. This early time point was chosen to better identify transcriptional patterns that may cause later functional and morphological changes.

An initial analysis of viral gene reads showed that, at 37°C, viral expression was clearly detectable at D2 for the D614G, Delta and Omicron BA.1 variants (Fig. 5A, top). The pattern of viral gene expression was comparable for the 3 variants, with N transcripts being the most abundant, followed by those coding for ORF1a and ORF1ab. The percentage of viral reads tended to be higher for BA.1 compared to D614G and Delta (Fig. 5B, top). It was notable that the percentage of viral reads reached above 30% of total mapped reads for epithelia infected by BA.1, highlighting how SARS-CoV-2 could usurp cellular resources for its own replication. The analysis of epithelia collected after 2 days of infection at 33°C showed a contrasting situation (Fig. 5A-B, bottom), with minimal detection of viral reads for the D614G and Delta variants, while the extent of BA.1 replication was high (26% of total mapped reads) and close to that observed at 37°C. The RNAseq analysis thus confirmed the replicative advantage of the BA.1 variant at nasal physiological temperature.

**Figure 5:**
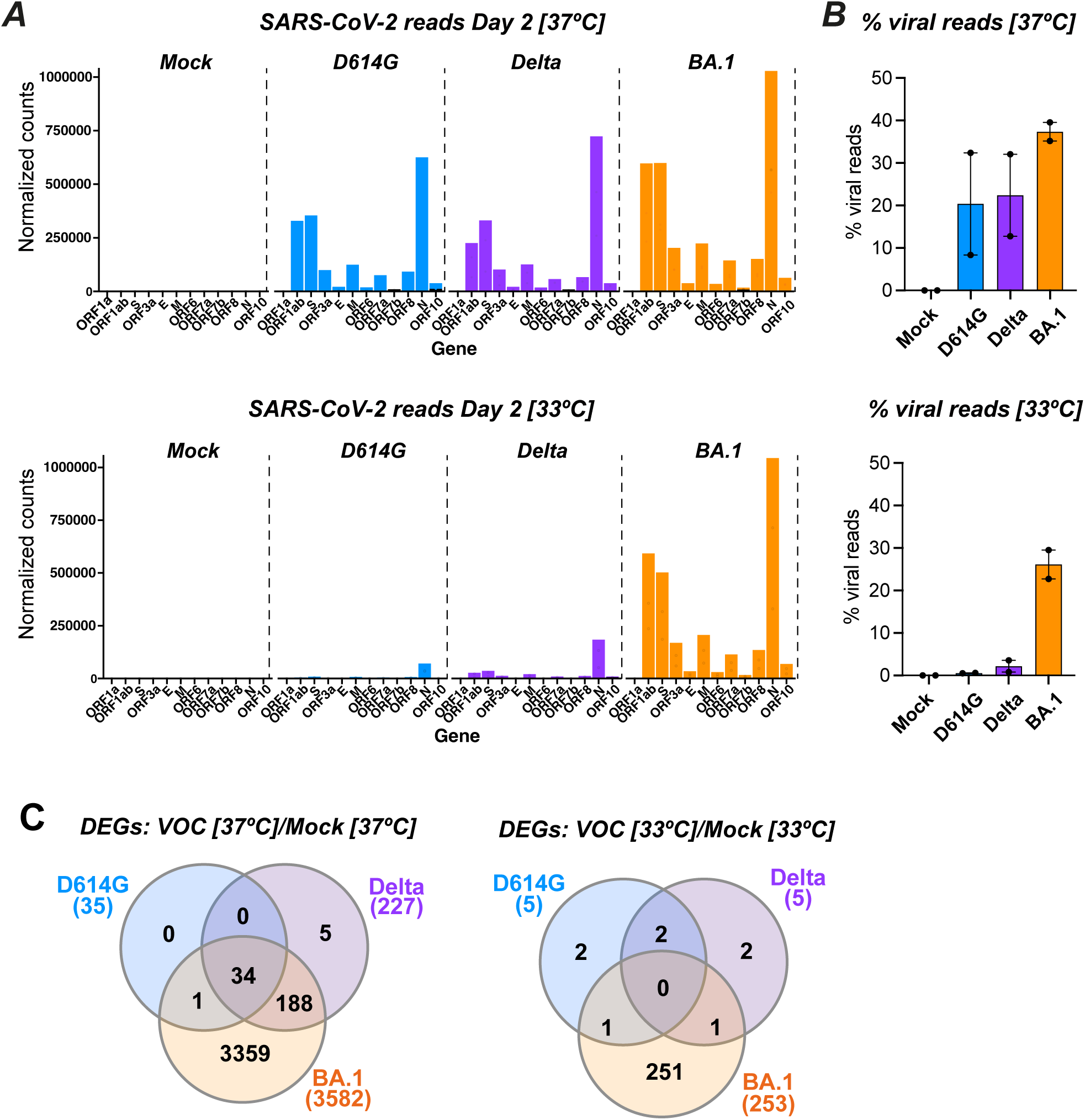
Temperature-dependent alteration of the cellular transcriptomic profile upon SARS-CoV-2 infection. A bulk RNA-seq analysis was performed at 2 dpi on reconstructed nasal epithelia either mock-infected or infected with SARS-CoV-2 variants, using cultures grown at 37°C or 33°C. (A) Distribution of SARS-CoV-2 reads at 37°C (top graph) and 33°C (bottom graph), showing the mean normalized counts per million for the different SARS-CoV-2 transcripts (n=2 replicates). (B) The percentage of viral reads among total mapped reads (cellular and viral) are reported for n=2 replicate samples infected at 37°C (top) and 33°C (bottom). The median and 95% confidence intervals are reported. (C) Venn-diagram showing the number of differentially expressed cellular genes (DEGs) shared between the different infected sample at 37°C (left graph) and 33°C (right graph). DEGs were determined with the DESeq2 package, with mock-infected samples used as a reference. The total number of DEGs for each variant is reported below the variant name.

Differentially expressed genes (DEGs) were defined as cellular genes that showed at least a 2-fold change in expression upon infection with an adjusted P value <0.05. The number of DEGs depended both on the nature of the infecting viral variant and on temperature (Fig. 5C). At 37°C, transcriptional dysregulation was low for D614G (n=35 DEGs), moderate for Delta (n=227), and marked for BA.1 (n=3582). Thus, the impact of infection on cellular transcripts was strongly variant-dependent, while viral transcripts at the same time point differed only to a moderate extent between variants. The cumulative effects of slight differences in viral replication kinetics may contribute to this variable effect on cellular transcriptomes. At 33°C, the D614G and Delta variants induced only minimal changes in cellular gene expression (n=5 DEGs for both viruses), while BA.1 caused clear cellular gene dysregulation (n= 253 DEGs), that remained however lower than that observed at higher temperature (Fig. 5C).

Interestingly, functional enrichment analysis revealed that the biological processes dysregulated by infection at D2 depended on the viral variant considered (Sup. Fig. S3). At 37°C, the top two gene ontology (GO) terms associated to transcriptional changes were “defense response” to viruses and symbionts for D614G and Delta, while the top two GO terms were “cilium organization” and “cilium assembly” for BA.1. At the lower temperature of 33°C, the same cilium-related GO terms were the most dysregulated for BA.1, while changes in biological processes were minimal for D614G and Delta (counts ≤3). Based on these findings, we analyzed more in depth the transcriptional changes for genes involved in cilia structure and function (Fig. 6A). In a volcano plot analysis, we compared the expression of genes specific for ciliated cells and goblet cells, as a balance between these two cell types had been previously reported in airway epithelia [36]. The analysis confirmed that Omicron BA.1 infection at 37°C induced a marked downregulation in the expression of ciliated cell genes, with a concomitant upregulation in the expression of goblet cell genes (Fig. 6A, right plot). In contrast, Delta infection caused only a slight upregulation of goblet cell gene expression, and D614G infection did not modulate ciliated nor goblet cell specific genes at this early D2 time point (Fig. 6A, middle and left plots). The same analysis carried out on epithelial samples infected at 33°C for 2 days confirmed the decrease ciliated cell gene expression and the increase in goblet cell gene expression induced by Omicron BA.1, while changes were not detected in these two gene sets upon D614G or Delta infection (Sup. Fig. S4A). Thus, Omicron BA.1 induced early D2 changes in transcriptional patterns that could account for the loss of motile cilia observed morphologically at the later D4 stage. The slower replication kinetics of D614G and Delta may explain the limited extent of transcriptional changes observed for these two variants at D2.

**Figure 6:**
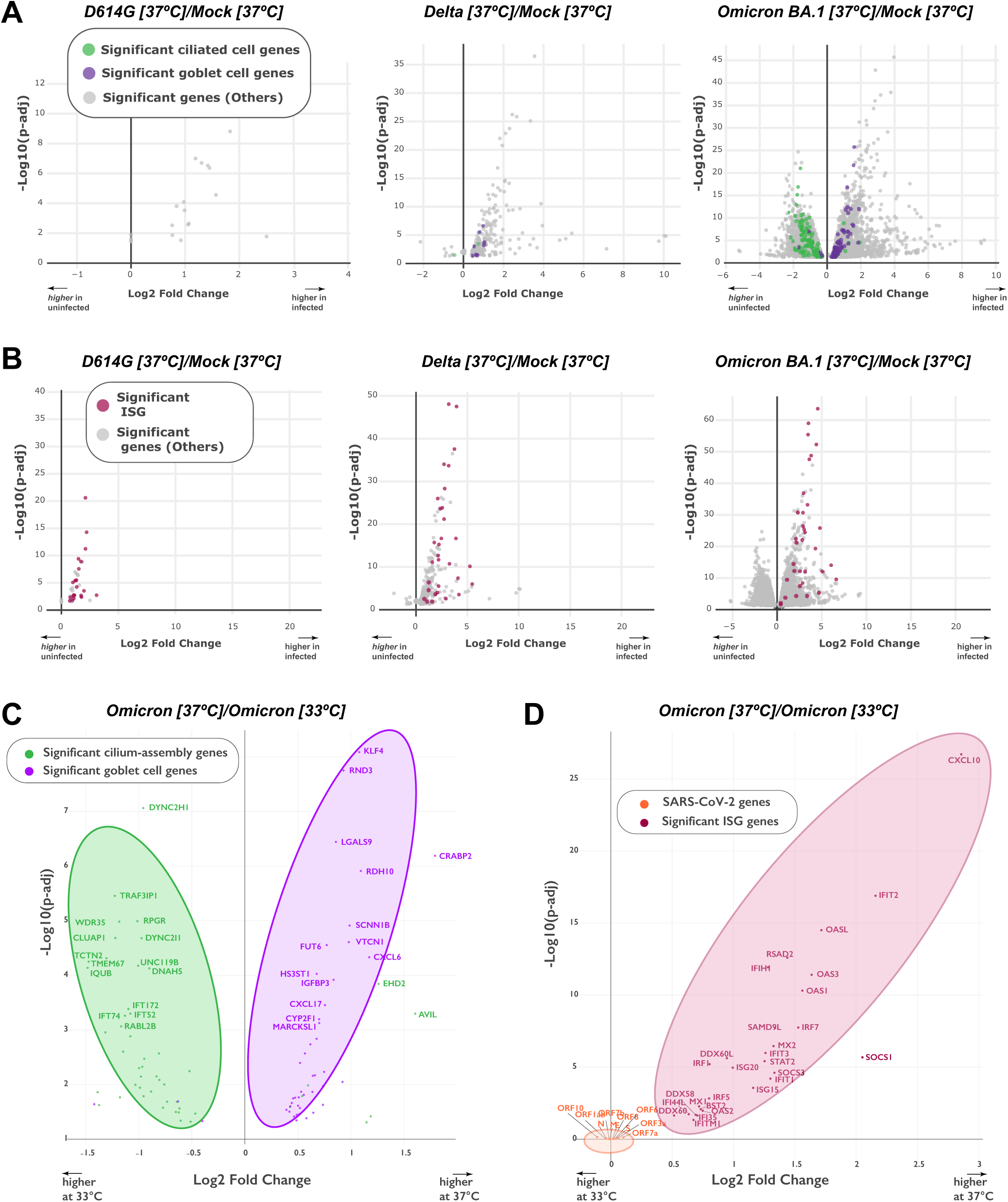
Transcriptional downregulation of ciliary genes and upregulation of IFN-stimulated genes (ISG) are variant- and temperature-dependent. (A) Volcano-plots depicting the differential regulation of ciliated cell genes (green) and goblet cell genes (purple) at 37°C in variant-infected samples compared to mock-infected samples at 2 dpi. Ciliated cell and goblet cell genes were defined by the clusterProfiler package. The log2 fold change in gene expression is shown on the x axis and the adjusted p value on the y axis. Genes overexpressed in infected samples are distributed to the right on the x axis. Significant DEGs that do not belong to the ciliated cell or goblet cell category are represented in grey. (B) Volcano-plots depicting the differential regulation of ISGs (red) and viral genes (orange) at 37°C in variant-infected samples compared to mock-infected samples at 2 dpi. The representation is similar to that in (A). (C, D) Volcano plots showing the impact of temperature on the differential expression of ciliated and goblet cell genes (C) or of ISGs and viral genes (D) in Omicron BA.1 infected samples at 2 dpi. Genes that are preferentially expressed at 37°C are distributed to the right on the x axis. Genes symbols are reported for the most dysregulated cellular genes and all viral genes.

We then performed a volcano plot analysis for interferon-stimulated genes (ISG), which represent the major gene set modulated early during antiviral defense (Fig. 6B). Upon infection at 37°C, the three variants did induce ISG transcripts at D2, with a more marked upregulation for the BA.1 and Delta variants compared to D614G (Fig. 6B). In contrast, at 33°C, the induction of ISG transcripts remained minimal for Omicron and undetectable for the Delta and D614G variants (Sup. Fig. S4B). To further explore this temperature-dependent effect, we directly compared gene expression patterns induced by Omicron infection at 33°C and 37°C (Fig. 6C-D). These comparisons were not performed for D614G and Delta, which showed too few DEGs at 33°C. Omicron BA.1 was found to more efficiently downregulate ciliated cell genes and upregulate goblet cell genes at 37°C than 33°C (Fig. 6C), consistent with the trend for a higher motile cilia loss observed at 37°C by immunofluorescence (Fig. 4). The most significantly downregulated cilia genes observed at 37°C coded for *DYNC2H1*, a microtubule-associated dynein; *TRAF3IP1*, a microtubule-interacting protein associated with *TRAF3*; *RPGR*, a guanyl nucleotide exchange factor involved in ciliation; and *WDR*, a component of the intraflagellar transport complex. Thus, both structural and regulatory components of motile cilia were affected by Omicron infection.

Analyzing temperature effects on antiviral defense gene expression showed that Omicron BA.1 infection induced a markedly stronger upregulation of ISGs at 37°C than at 33°C (Fig. 6D). The most significantly upregulated ISG at 37°C included the chemokine *CXCL10*, which attracts immune cellular effectors; the *IFIT2* antiviral protein, which inhibits the expression of uncapped viral RNA; the *OASL* RNA binding protein, which potentiates IFN production; and *RSAD2* or viperin, an enzyme that produces an antiviral ribonucleotide. Multiple mechanisms targeting RNA virus replication were thus induced by Omicron at 37°C. Viral transcripts were represented on the same volcano plot, to evaluate the association between the triggering factors (viral RNA) and the epithelial antiviral response (ISG expression). Interestingly, while the difference in ISG expression was marked at 37°C vs 33°C, the difference in BA.1 viral expression was minimal. Indeed, viral transcripts were grouped close to the origin of the volcano plot, pointing to a lack of significant changes in viral expression when comparing the two temperatures (Fig. 6D, orange symbols). These findings highlight the capacity of Omicron BA.1 to replicate efficiently at nasal physiological temperature while inducing only a limited antiviral defense reaction in the epithelium.

### Limited induction of IFNs by Omicron at nasal physiological temperature

To further evaluate the epithelial antiviral response upon SARS-CoV-2 infection, we directly measured the secretion of IFNs in apical culture supernatants at D2 and D4 (Fig. 7). Using high-sensitivity immunoassays [37][32], we detected an induction of both type I (IFN-β) and type III (IFN-λ1 and IFN-λ2/3) IFNs between D2 and D4 at 37°C, with a positive trend for all the variants tested (Fig. 7A, top row). Of note, the Delta variant induced the strongest IFN induction at D4, while IFN concentrations observed for Omicron BA.1 appeared intermediate, and those observed for the other variants Alpha and D614G remained low (IFNλ1 induction between D2 and D4: P<0.05 for Delta and for BA.1. IFNλ2/3 induction between D2 and D4: P<0.01 for Delta; P<0.05 for BA.1). A similar phenomenon was observed in epithelial cultures infected at 33°C (Fig. 7A, bottom row), with a more efficient induction of type I and type III IFNs by Delta compared to the other variants tested, including Omicron. Indeed, Delta was the only variant to induce a detectable production of IFN-β at physiological nasal temperature.

**Figure 7:**
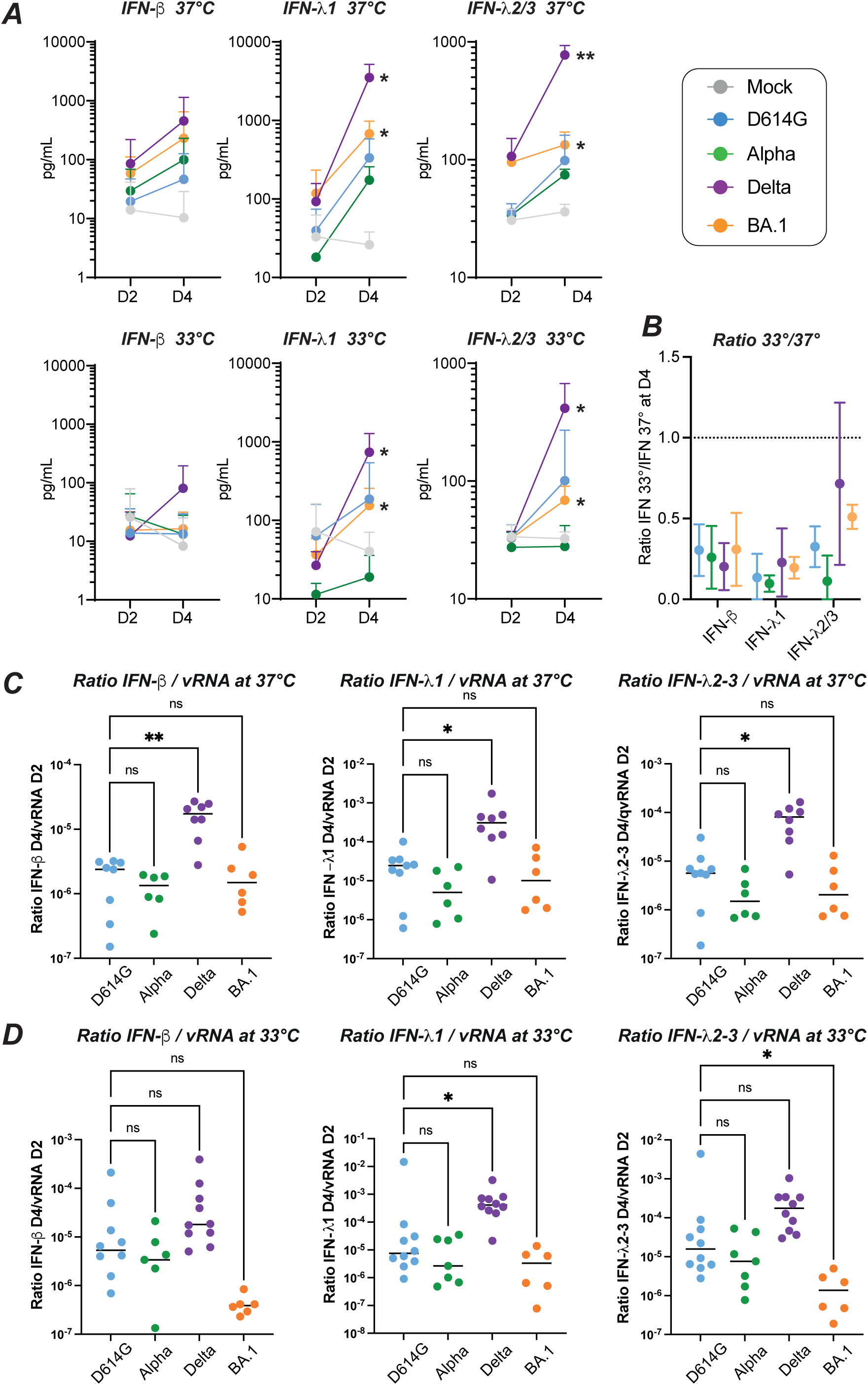
Differential induction of type I/III IFNs by SARS-CoV-2 variants. (A) The induction of IFN-β (left), IFN-λ1 (middle) and IFN-λ2/3 (right) release in apical supernatants of reconstructed nasal epithelia infected at 37°C (top panels) and 33°C (bottom panels) was measured by digital ELISA (Simoa) and Legendplex assays. The mean +/- SD are reported for n=3 to 6 independent biological replicates. The induction of IFN between 2 dpi (D2) and 4 dpi (D4) was evaluated with a paired t-test. (B) The ratio of IFN measured at 33°C to that measured at 37°C is reported for the 4 variants tested (colored symbols) and the 3 IFNs measured (IFN-β, left; IFN-λ1, middle; IFN-λ2/3, right) at the D4 time point. The mean +/- SD are reported for n=4 to 5 independent biological replicates. The dashed line represents a ratio value of 1. (C) The ratio of IFN released at 4 dpi to the viral RNA released at 2 dpi is reported at 37°C (C) and 33°C (D) for IFN-β (left plots), IFN-λ1 (middle plots) and IFN-λ2/3 (right plots). Medians are reported. Statistical comparisons were done between the D614G reference variants and all the other variants tested, using the Kruskal-Wallis test with Dunn’s correction; ns: not significant; * P<0.05; ** P<0.01.

To compare the production of IFNs at the two temperatures tested, we computed the ratio of concentrations measured at 33°C to those measured at 37°C for the D4 timepoint (Fig. 7B). The production of IFNs proved consistently weaker at 33°C, as indicated by ratios below 1 for all the IFNs tested. For type III IFNs, the ratios tended to be higher for Delta and Omicron compared to Alpha. However, taken together, all the SARS-CoV-2 variants tested triggered lower antiviral responses at nasal physiological temperature.

An inter-variant comparison of apical IFN concentrations measured at D4 at 37°C (Sup. Fig. S5A) confirmed that Delta was the only variant to induce a significantly higher IFN secretion compared to the reference variant D614G, independently of the IFN tested (P<0.01 for IFN-β, IFN-λ1 and IFN-λ2/3). Omicron induced an intermediate IFN secretion that did not differ significantly from that measured for D614G. A similar pattern was observed at 33°C (Sup. Fig. S5B). To determine whether these findings applied to other IFNs, we measured the secretion of IFN-α at D4 in a subset of experiments (Sup. Fig. S5C). We observed a trend for a higher induction of IFN-α by Delta compared to Omicron at 37°C, consistent with findings obtained for other type I and type III IFNs. In contrast, no induction of IFN-α could be detected at 33°C for all the variants tested, suggesting that IFN-α induction was highly temperature-dependent.

It was intriguing that Omicron BA.1 induced only a moderate induction of IFNs at 33°C, while this variant had an early replicative advantage at this temperature. To further explore this point, we computed the ratio of IFN secreted at D4 reported to the number of inducing viral RNA copies measured at the early D2 time point. At 37°C, the Delta variant showed significantly higher IFN to viral RNA ratios compared to the D614G reference, while Omicron BA.1 showed ratios equivalent to those of D614G (Fig. 7C). Interestingly, at 33°C, Omicron BA.1 showed lower IFN to viral RNA ratios compared to D614G (P<0.05 for IFNλ2/3), while a trend for higher ratios was maintained for Delta (P<0.05 for IFNλ1). This analysis confirmed that Delta was a potent inducer of type I and type III IFNs. In contrast, Omicron BA.1 was characterized by a weak induction of IFNs relative to its replication capacity, especially at nasal physiological temperature.

## DISCUSSION

This study highlights that the SARS-CoV-2 Omicron variant evolved several traits that converged in facilitating viral spread. First, Omicron displayed an early replicative advantage in primary human nasal epithelial cells, which represent the main source of transmissible virus. Second, the increase in Omicron replication was more notable at 33°C, the temperature typically present in human nasal passages, resulting in a physiologically relevant replicative advantage. Third, while Omicron did not entirely disrupt the epithelial structure when replicating at 33°C, it did cause measurable decreases in epithelial integrity and in motile cilia coverage. These changes are likely to perturb the mechanism of mucociliary clearance, as we reported previously [32], and could thus facilitate the spread of viral particles within the airways. Fourth, at this physiological temperature, Omicron remained a moderate inducer of IFN responses relative to its replicative capacity in the nasal epithelium. Considering the key role of the type III IFN response in limiting the early spread of viruses at epithelial surfaces [38][39], it is relevant that Omicron induced comparatively less IFN-λ1 and IFN-λ2/3 than the Delta variant. The combined advantages of increased viral replication, perturbation of motile cilia, and escape from the innate antiviral response in the nasal epithelium likely contributed to the worldwide spread of the original Omicron BA.1 variant and to the continued dominance of Omicron-derived variants since 2022. While escape from the adaptive immune response clearly played an important role in Omicron emergence [9][8], our findings argue for an additional role of intrinsic viral properties in the evolutionary success of the Omicron variant.

The IFN response is a key determinant of Covid severity, as shown by the association of inborn errors in the type I IFN pathway and critical Covid pneumonia [40]. The importance of the IFN pathway is also supported by the detection of auto-antibodies capable of neutralizing type I IFNs in up to 20% of patients who had fatal Covid [41]. A decreased IFN response could be directly observed in the nasopharyngeal swabs from patients with severe Covid [36]. In face of the major selective pressure exerted by the IFN response, SARS-CoV-2 has evolved a variety of countermeasures, with multiple viral genes involved in dampening the induction of IFNs and/or inhibiting the function of ISGs [42][43]. Indeed, SARS-CoV-2 was shown to be more efficient at escaping the innate antiviral response than common cold coronaviruses, helping to explain why the former is more pathogenic [44].

Of interest, successive SARS-CoV-2 variants have shown an evolution in their capacity to escape the innate antiviral response. The early variant Alpha was shown to induce lower levels of IFNs than the original SARS-CoV-2 strain, due to increased expression of viral genes involved in innate immune antagonism, such as ORF6, ORF9b, and a truncated form of the nucleocapsid [45][46]. Studies of later variants have yielded variable findings, with some studies showing an increasing capacity to escape the antiviral effects of type I IFNs from Alpha to Delta to Omicron [29], others showing a better escape for Alpha and Omicron compared to Delta [30][47], and still others reporting a more efficient escape for Delta compared to Omicron BA.1 [48]. One reason for these divergent findings could be that some studies were performed on transformed cell lines such as Calu-3, in which Omicron shows an attenuated phenotype, in contrast to the fast-replicating phenotype observed in primary epithelial cells. Another confounding factor may be the procedure used to titrate viral stocks, which often relies on serial dilutions in transformed cells lines such as Vero-E6-derived cells. As Omicron shows an attenuated phenotype in such cell lines, the resulting viral stocks tend to be underestimated in terms of infectivity for primary cells. To avoid these caveats, we chose to normalize the viral stocks based on viral RNA content, which is a trigger of the IFN response [49]. Further, we chose to analyze viral replication kinetics and IFN induction in a human primary nasal cell-based model, which is more likely to yield information relevant to human physiology.

Based on this strategy, we observed that the induction of type I and type III IFNs in primary nasal epithelial cells was low for Alpha, intermediate for Omicron, and significantly higher for Delta. To consider the faster replication kinetics of Omicron, and hence the more rapid accumulation of viral RNA acting as a PAMP (Pathogen Associated Molecular Pattern), we computed the ratio of produced IFN to early viral RNA copies. This analysis pointed to a relatively low intensity of the IFN response to Alpha and Omicron, compared to a significantly higher induction of type I and III IFN for Delta. A possible reason for the strong innate response to Delta maybe the highly fusogenic phenotype of this variant [5][35], as fusion is per se a triggering event for the IFN response [50]. Fused syncytia are thought to have short life spans and to release DAMPs (Damage-Associated Molecular Patterns) when dying. It is thus relevant that we observed the formation of larger syncytia in the Delta infected cultures, consistent with a highly fusogenic phenotype for this variant. A limitation of our study is that the reconstructed epithelium model used is devoid of immune cells, so that we cannot evaluate the contribution of resident and inflammatory leukocytes to the mucosal IFN response. It is interesting, however, that an *in vivo* study of the innate response in nasopharyngeal swabs obtained during acute Covid found an inverse correlation between nasal ISG expression and Covid severity in patients infected with the ancestral and Delta variants, but not with the Omicron variant [51]. These *in vivo* findings are compatible with a stronger impact of the IFN response on Delta than on Omicron infections, consistent with our findings in primary nasal epithelial cells. Also relevant is a recent study showing that later Omicron variants, including BA.4 and BA.5, have further increased their suppression of innate immunity compared to BA.1 and BA.2, pointing to a continued evolution of SARS-CoV-2 variants towards improved innate immune evasion [31].

All the variants tested could replicate at both 33°C and 37°C, consistent with the dual tropism of SARS-CoV-2 for the upper and lower respiratory tract [52][53][54]. This contrasts with the preferential replication of common cold coronaviruses at 33°C, in line with their restriction to the upper respiratory tract [55]. At the early D2 time point, the Omicron variant had already reached its replication plateau when grown at 33°C, while the induction of ISGs proved minimal or undetectable at this temperature. Furthermore, at D2, the production of type I and type III IFNs was not induced in Omicron-infected cultures grown at 33°C. These observations fit with the long-held notion that the IFN response is inefficient and delayed at lower temperature, mostly due to limited transcription of ISGs and IFN genes [56][57]. Our data quantifying IFN-β and IFN-λ suggest that this temperature effect may also impact IFN protein secretion. It was interesting that Omicron showed a replicative advantage over other SARS-CoV-2 variants particularly in the low temperature condition. This observation suggests that Omicron is not only efficient at evading the IFN response, but that it possesses other fitness enhancing properties independently of its innate response evasion capacity.

Structural features may contribute to Omicron’s fitness at lower temperature, as its Spike protein was shown to have better thermostability, with a tighter packing of the 3 receptor-binding domains in their down state [58] [59]. The viral attachment step is also likely involved, as Omicron was proposed to have a higher affinity for its receptor ACE-2 [23][24] and a higher capacity to bind the plasma membrane of its target cells [25][26]. The SARS-CoV-2 fusion step depends on lipid membrane reorganization and is known to be temperature dependent, with a lower efficiency at 33°C than at 37°C [60]. As the Omicron Spike was proposed by some authors to switch more easily to a fusogenic conformation, it is possible that Omicron better overcomes the temperature-dependent barrier to fusion [25]. On the other hand, a more rapid switch to a fusogenic conformation may lead to viral inactivation by fusion with extracellular vesicles, which can act as traps for incoming virions [61]. It is thus interesting that the release of extracellular vesicles is decreased at lower temperature, which may facilitate the spread of fusion-prone variants at 33°C. The nature of the Omicron entry pathway remains to be precisely defined, as several groups have reported increased Omicron entry through a cathepsin-dependent endosomal pathway [27][28], while others have highlighted an increased dependence of Omicron entry on metalloproteases [62][14], and still others report a continued dependence on the surface protease TMPRSS2 to trigger Omicron Spike fusion [63]. Further studies are thus warranted to precisely characterize the post-attachment steps involved in Omicron entry in primary cells and determine whether they contribute to the replicative advantage of this variant at nasal physiological temperature.

Of note, Omicron perturbed the integrity of the motile cilia layer more efficiently than other variants. The differences were marked at lower temperature, with a significant decrease in TEER values compared to the reference D614G variant, indicative of increased epithelial permeability. In addition, Omicron induced more cell death and increased cilia loss at 33°C compared to other variants. Motile cilia loss can have several consequences on viral dissemination. At the cellular level, it can facilitate the dissemination of viral particles, which are released at the plasma membrane and will not have to cross a layer of packed cilia to reach neighboring cells and/or the mucus layer. The loss of motile cilia may also facilitate the passage of actin protrusions that push recently released SARS-CoV-2 virions towards the mucus layer [26]. At the epithelium level, the loss of motile cilia inhibits mucociliary clearance, as viral particles become trapped in a mucus layer that is not propelled anymore by ciliary beating [32]. This may locally limit the spread of viral particles to neighboring ciliated cells, as viral particles become less motile [64]. On the other hand, at the organ level, the loss of mucociliary clearance prevents the removal of viral particles trapped in mucus and their redirection towards the pharynx [65][66]. The loss of this clearance mechanism likely facilitates the viral invasion of epithelia situated lower in the respiratory tract [67], and thus promotes viral dissemination within the infected organism. Omicron outbreaks in Asia have been associated with a high rate of bacterial and fungal superinfections, which may also reflect an impairment of mucociliary clearance [68][69]. It is interesting that the Omicron variant can perturb mucociliary clearance at the temperature found in nasal passages, suggesting that this property is physiologically relevant and beneficial to the evolutionary fitness of this variant.

In conclusion, the present study highlights multiple features that likely contributed to the efficient transmission and worldwide dissemination of Omicron. This SARS-CoV-2 variant appears particularly adapted to the upper respiratory tract, based on its increased replicative capacity at the temperature found in nasal passages, its capacity to perturb the motile cilia layer, and its limited induction of the IFN epithelial response. Further development of 3D-tissue models that include innate and adaptive immune cells should help achieve an integrated view of the parameters that control SARS-CoV-2 variant dissemination in the upper airways.

## MATERIALS AND METHODS

### SARS-CoV-2 infection of reconstructed human nasal epithelia

Reconstructed human nasal epithelial cultures (MucilAir^TM^) differentiated *in vitro* for at least 4 weeks were purchased from Epithelix (Saint Julien-en-Genevois, France). Cultures were maintained in air/liquid interface (ALI) conditions in transwells, with 700 μL of MucilAir^TM^ medium (Epithelix) in the basal compartment and kept at 37 °C under a 5% CO2 atmosphere. For SARS-CoV-2 infection, the apical side of ALI cultures was washed 20 min at 37°C in Mucilair^TM^ medium to remove mucus. Cells were then incubated with the equivalent of 10^6^ RNA viral copies/µL of viral stock for each of the SARS-CoV-2 variants tested: Wuhan, D614G, Alpha, Beta, Gamma, Delta and Omicron BA.1. The source of the viral isolates used is reported in Supplementary Table 1. The viral input was diluted in DMEM medium to a final volume 100 μL and left on the apical side for 4 h at 37°C. Control wells were mock-treated with DMEM medium (Gibco) for the same duration. Viral inputs were removed by washing three times with 200 μL of PBS (5 min at 37°C) and once with 200 μL MucilAir^TM^ medium (20 min at 37 °C). The basal medium was replaced every 2–3 days. Apically released viruses were collected by incubation with 200 μL MucilAir^TM^ medium (20 min at 37°C) at days 1, 2, and 4 post-infection. The integrity of the epithelia was monitored by measuring the trans-epithelial electrical resistance (TEER) at the same days. To do so, after apical wash, the transwells were transferred into a new 24-well plate and DMEM medium was added to both the apical (200 μL) and basal (700 μL) sides. The TEER was then measured using an Evom3 ohmmeter (World Precision Instruments). At day 4, cultures were fixed in paraformaldehyde 4% for 30 min, washed twice with PBS, and stored in PBS at 4°C until immunofluorescence staining.

### Viral RNA quantification

Culture apical supernatants collected at days 1, 2 and 4 post-infection were inactivated for 20 min at 80°C and then subjected to RT-qPCR analysis, using a Luna Universal One-Step RT-qPCR Kit (New England Biolabs), following the manufacturer’s instructions. SARS-CoV-2 RNA was quantified in a final volume of 5 μL per reaction in 384-well plates using SARS-CoV-2 N-specific primers (Forward 5’-CGA AGG TGT GAC TTC CAT G-3’; Reverse 5’- TAA TCA GAC AAG GAA CTG ATT A-3’) on a QuantStudio 6 Flex thermocycler (Applied Biosystems). A standard curve was performed in parallel using purified SARS-CoV-2 viral RNA (EURM019, Sigma Aldrich).

### Immunofluorescence labeling

Membranes supporting the fixed MucilAir^TM^ cultures were cut in four using a scalpel blade. Membrane pieces were placed in 10 μL drops of phosphate buffer saline (PBS) onto parafilm and then permeabilized in PBS 0.5% Triton-X100 for 20 min at room temperature (RT). Samples were blocked in PBS - 0.1% Tween - 1% bovine serum albumin (BSA) - 10% fetal bovine serum - 0.3 M glycine for 30 min at RT. Samples were then incubated overnight at 4°C with the following antibodies: pan-SARS-CoV-2 anti-Spike mAb102 human monoclonal antibody [70] at 1 μg/mL; rabbit anti-cleaved caspase 3 primary antibody (9661, Cell Signaling; 1:100 dilution); AF647-conjugated rabbit anti-alpha acetylated tubulin (ab218591, Abcam; 1:100 dilution). After washing, a secondary anti-human IgG antibody conjugated to AF555 (A-21422, Thermo Fisher Scientific) was added at 1:400 for 1 hour at RT. Primary and secondary antibodies were diluted in PBS - 0.1% Tween - 1% BSA. Samples were counterstained with Hoechst and mounted in FluoromountG (Thermo Fisher Scientific) before observation with an LSM 710 confocal microscope (Zeiss) or with a BC43 spinning disk confocal microscope (Andor).

### LDH cytotoxicity assay

Diluted apical culture supernatants (1:25) were pretreated with Triton-X100 1% for 2 h at room temperature for viral inactivation. Lactate dehydrogenase (LDH) dosage was performed using the LDH-Glo™ Cytotoxicity Assay kit (Promega) following the manufacturer’s instructions. Luminescence was measured using an EnSpire luminometer (Perkin Elmer).

### Cytokine measurements

Apical culture supernatants were pretreated with Triton-X100 1% (final concentration) for 2 h at room temperature for viral inactivation. IFN-β protein was quantified using Simoa digital ELISA technology as previously described [32][37]. Alternatively, IFN-β and IFN-α were quantified simultaneously in a Simoa multiplexed assay. Briefly, inactivated supernatant were diluted 1:10 in Diluent B (Quanterix, Billerica, MA, USA) containing 1% Triton-X100 (final concentration). IFN-α and IFN-β concentrations were quantified with a Simoa assay developed on an HD-X instrument with Quanterix Homebrew kits. The assay configuration was a 2-step triplex associating IFN-α, IFN-β and IFN-λ3 (data non shown), using 150pM SBG and 50uL RGP reagents. The IFN-α plex quantifies all IFN-α subtypes with a similar sensitivity, as previously reported [37], with results expressed in IFN-α17 equivalents. The antibodies used were cloned from APECED/APS1 patients: The 8H1 antibody clone was used as a capture antibody after coating on paramagnetic beads (0.3 mg/mL); the 12H5 clone was biotinylated (biotin/antibody ratio = 30:1) and used as the detector at a concentration of 0.3 µg/mL. The recombinant human IFN-α17 protein (#11150, PBL Assay Science, Piscataway, NJ, USA) was used as reference to quantify concentrations. The limit of detection was between 0.59-1.43 fg/mL equivalent IFN-α17, including the dilution factor. The IFN-β plex assay is described more in details in [71]. Briefly, The 710906-9 IgG1-κ mouse monoclonal antibody (PBL Assay Science) was used as a capture antibody to coat paramagnetic beads (0.3 mg/mL); the 710323-9 IgG1-κ mouse monoclonal antibody (PBL Assay Science) was biotinylated (biotin/antibody ratio = 40/1) and used as the detector antibody at a concentration of 0,3 µg/mL. The recombinant IFN-β protein (#11415, PBL Assay Science) was used as reference to quantify concentrations. The limit of detection was between 0.01-0.03 pg/mL IFN-β, including the dilution factor.

#### Quantification of IFN-λ1 and IFN-λ2/3 proteins using Legendplex technology

On day of assaying, supernatants were centrifuged at 300g for 5 minutes. The concentration of 13 different analytes were measured using the LEGENDplex Human Anti-Virus Response Panel (13-plex) with V-bottom plate kit (cat. No. 740390) and the manufacturer protocol was followed (BioLegend, San Diego, California, USA). Plates were read on a BD Fortessa II 5-color flow cytometer (Franklin Lakes, New Jersey, USA), and the resulting MFI data were analyzed using the BioLegend portal to predict analyte concentration (https://legendplex.qognit.com/).

### Scanning electron microscopy

PFA-fixed epithelial samples were post-fixed with 2.5% glutaraldehyde in 0.1 M cacodylate buffer for 1h at RT. Samples were then washed in 0.1 M cacodylate buffer and several times in water, and then incubated in 1% osmium tetroxide for 1h. After dehydration by incubation in increasing concentrations of ethanol (35%, 70%, 85%, 95%, and 100%), samples were treated with hexamethyldisilazane for 10 min for chemical critical point drying. Specimens were then sputter-coated with up to 9 Å gold/palladium conductive layer using a gun ionic evaporator PEC 682. Images were acquired on a JEOL JSM 6700F field emission scanning electron microscope operated at 7 kV.

### RNAseq

#### Sample preparation and sequencing

After 2 days of infection, the epithelial cultures were washed in cold PBS and then lyzed in 150 μL of Trizol reagent (Thermofisher scientific) added to the apical side of the insert. RNAs were purified using the Direct-zol miniprep kit (ZR2080, Zymo Research). RNA integrity and concentration were verified by Bioanalyzer (Agilent Technologies). mRNA enrichment, strand specific library preparation, and paired-end sequencing (100 bp) on the DNBSEQ™ platform were performed at BGI (Poland).

#### Processing of raw sequencing data

Raw paired-end RNA-sequencing data obtained from BGI were analyzed using FastQC v0.11.9 for quality control [72] and trimmed with fastp v0.23.2 [73].Trimmed reads were mapped using HISAT2 v2.2.1 [74] to the following reference genomes: concatenated GRCh38 (human) genome and wuhCor1 (SARS-CoV-2) reference genomes obtained from the UCSC Genome Browser (http://hgdownload.soe.ucsc.edu/goldenPath). Duplicate alignments were labelled with the markDuplicates function from Picard v2.23.4 (http://broadinstitute.github.io/picard/). Marked alignment files were further processed using Sambamba 0.7.1 [75] keeping only uniquely mapping reads after duplicate removal.

#### Analysis of RNA-seq data

Mapped reads were quantified with featureCounts v2.0.0 using GRCh38 Gencode v36 annotation, following differential expression analysis with DESeq2 [76] with two replicates per condition. Genes were considered differentially expressed with an absolute log2 fold change value > 1 and adjusted p-value < 0.05. Functional enrichment analysis was performed using the clusterProfiler [77] and enrichR packages [78]. Data visualization, including volcano plots, was carried out using custom scripts with the ggplot2 R package [79]. To quantify the proportion of viral reads within each sample, the number of all mapped reads aligning to the SARS-CoV-2 genome was calculated using featureCounts v2.0.0 and further normalized by the total number of mapped reads per sample.

### Statistics and reproducibility

Statistical analyses were performed with the Prism software v8.4.3 (GraphPad) for all figures except for RNAseq analyses described above. The non-parametric Kruskal-Wallis test with Dunn’s correction for multiple testing was used in all cases, except for paired analyses, where a paired t-test was used. All tests were two-sided. Statistical significance was assigned when p values were <0.05. The nature of statistical tests and the number of experiments (n) are reported in the figure legends.

### Data availability

The data underlying this article are available in NCBI Gene Expression Omnibus (GEO) at https://www.ncbi.nlm.nih.gov/geo/ and can be accessed with the GSE295759 number.

## ACKNOWLEDGMENTS

We thank Hugo Mouquet and Cyril Planchais (Humoral Immunity Unit, Institut Pasteur) for providing the Spike-specific antibody. We acknowledge the Photonic BioImaging UtechS, supported by the French National Research Agency (France BioImaging; ANR-10–INSB–04; Investments for the Future) for help with image acquisition.

This work was supported by the 3D-LUNGO project (reference ANRS-23-PEPR-MIE 0001) supported by the program France 2030 and managed by the French National Agency for AIDS and Emerging Diseases Research ANRS-MIE (to LAC and SG); the COROCHIP project funded by the Institut Pasteur COVID-19 RP call (to LAC and SG); the Tous Unis COVID project (reference PR-166156) funded by the Fondation de France (to LAC); the PFR7 project funded by the Urgence COVID-19 Fundraising Campaign of Institut Pasteur (to LAC); the PERSICOT project (reference ECTZ213626) funded by ANRS-MIE (to LAC); the Steroid Response project funded by the Institut Pasteur COVID-19 RP call (to DD). RR was the recipient of a Sidaction fellowship, SL of an ANRS-MIE / Fondation pour la Recherche Médicale (FRM) fellowship, and CC from a France 2030 / ANRS-MIE PEPR fellowship.

## AUTHOR CONTRIBUTIONS

Conceptualization: BFF RR VM OS DD TM SG LAC

Investigations: BFF, RR, VM, SL, CC, RJM, RV, VB, EM, FGB

Funding acquisition: LAC, SG, OS, TM, DD

Formal analysis: BFF, AM, RV, TM

Writing - Original draft preparation: BFF, LAC

Writing - Review & Editing: all the authors

## COMPETING INTERESTS

The authors declare no competing interests.

**Supplementary Figure S1: Temperature-dependent replication of the Beta and Gamma SARS-CoV-2 variants in reconstructed human nasal epithelia.** The viral load in apical supernatants was quantified by RT-qPCR for samples infected by the following SARS-CoV-2 lineages: D614G, Beta, and Gamma. (A, B) Kinetics of viral replication at 37°C (A) and at 33°C (B), with means and SD reported (n=4 to 14 independent samples per point). (C,D) Comparison of apical viral loads measured at 2 days post infection (dpi) at 37°C (C) and 33°C (D). Medians are reported. (E) The ratio of viral RNA copies measured at 33°C to those measured at 37°C is shown at different dpi, with means and SD reported (n=4 to 14 independent samples per point).(C, D, E) Statistical comparisons were done between the D614G reference and all other variants or the mock condition, using the Kruskal-Wallis test with Dunn’s correction; ns: not significant; **** P<0.0001.

**Supplementary Figure S2: Infection by SARS-CoV-2 variants induce a loss of motile cilia.** Scanning electron microscopy images of reconstructed nasal epithelia at 4 days post-infection. The epithelia were infected at 37°C (left) or at 33°C (right) with the D614G, Delta or Omicron, BA.1 variant (shown in rows 2, 3, 4, respectively) or were mock-infected (first row). The scale bar represents 10 µm. Red symbols: * membrane protrusion surrounding cilia; ** blebbing cell.

**Supplementary Figure S3: Functional enrichment analysis of transcriptomes from reconstructed nasal epithelia at day 2 post-infection.** The significantly dysregulated gene ontology (GO) terms are reported for epithelial samples infected at 33°C (top) or 37°C (bottom) with the variants D614G (left), delta (middle), and Omicron BA.1 (right). The number of differentially expressed genes (DEGs) belonging to each GO term is reported on the x axis. The size and color of symbols correspond to the gene ratio and p-value for each GO term, respectively. Among the 5 top dysregulated GO terms, those corresponding to “cilium organization” and “defense response” are highlighted in orange and yellow, respectively.

**Supplementary Figure S4: Limited changes in transcriptional profiles of reconstructed nasal epithelia infected at 33°C.** (A) Volcano-plots depicting the differential regulation of ciliated cell genes (green) and goblet cell genes (purple) at 33°C in variant-infected samples compared to mock-infected samples at 2 dpi. The log2 fold change in gene expression is shown on the x axis and the adjusted p value on the y axis. Genes overexpressed in infected samples are distributed to the right on the x axis. Significant DEGs that do not belong to the ciliated cell or goblet cell category are represented in grey. (B) Volcano-plots depicting the differential regulation of ISGs (red) and viral genes (orange) at 33°C in variant-infected samples compared to mock-infected samples at 2 dpi. The representation is similar to that in (A).

**Supplementary Figure S5: Induction of type I and type III IFNs by SARS-CoV-2 variants at day 4 post-infection.** (A) The induction of IFN-β (left), IFN-λ1 (middle) and IFN-λ2/3 (right) release in apical supernatants of reconstructed nasal epithelia infected at 37°C was measured by digital ELISA (IFN-β) and Legendplex assays (IFN-λ). (B) Similar graphs as in (A), but for infections carried out at 33°C. (C) The induction of IFN-α release in apical supernatants of reconstructed nasal epithelia infected at 37°C (left) and 33°C (right) was measured by digital ELISA, with results expressed as IFN-α17 equivalent. (A-C) Medians are reported. Statistical comparisons were done between the D614G reference variants and all the other variants tested, using the Kruskal-Wallis test with Dunn’s correction; ns: not significant; ns: not significant; * P<0.05; ** P<0.01; ** P<0.001.

**Supplementary Table 1: Viral stocks used in the study.**

## Notes

### Competing Interest Statement

The authors have declared no competing interest.

## REFERENCES

1. Carabelli AM, Peacock TP, Thorne LG, Harvey WT, Hughes J, de Silva TI, et al. SARS-CoV-2 variant biology: immune escape, transmission and fitness. Nat Rev Microbiol. 2023;21: 162–177. doi:10.1038/s41579-022-00841-7

2. Yajima H, Nomai T, Okumura K, Maenaka K, The Genotype to Phenotype Japan (G2P-Japan) Consortium, Ito J, et al. Molecular and structural insights into SARS-CoV-2 evolution: from BA.2 to XBB subvariants. mBio. 2024;15: e03220–23. doi:10.1128/mbio.03220-23

3. Korber B, Fischer WM, Gnanakaran S, Yoon H, Theiler J, Abfalterer W, et al. Tracking Changes in SARS-CoV-2 Spike: Evidence that D614G Increases Infectivity of the COVID-19 Virus. Cell. 2020;182: 812–827.e19. doi:10.1016/j.cell.2020.06.043

4. Twohig KA, Nyberg T, Zaidi A, Thelwall S, Sinnathamby MA, Aliabadi S, et al. Hospital admission and emergency care attendance risk for SARS-CoV-2 delta (B.1.617.2) compared with alpha (B.1.1.7) variants of concern: a cohort study. The Lancet Infectious Diseases. 2022;22: 35–42. doi:10.1016/S1473-3099(21)00475-8

5. Mlcochova P, Kemp SA, Dhar MS, Papa G, Meng B, Ferreira IATM, et al. SARS-CoV-2 B.1.617.2 Delta variant replication and immune evasion. Nature. 2021;599: 114–119. doi:10.1038/s41586-021-03944-y

6. Saito A, Irie T, Suzuki R, Maemura T, Nasser H, Uriu K, et al. Enhanced fusogenicity and pathogenicity of SARS-CoV-2 Delta P681R mutation. Nature. 2022;602: 300–306. doi:10.1038/s41586-021-04266-9

7. Viana R, Moyo S, Amoako DG, Tegally H, Scheepers C, Althaus CL, et al. Rapid epidemic expansion of the SARS-CoV-2 Omicron variant in southern Africa. Nature. 2022;603: 679–686. doi:10.1038/s41586-022-04411-y

8. Planas D, Saunders N, Maes P, Guivel-Benhassine F, Planchais C, Buchrieser J, et al. Considerable escape of SARS-CoV-2 Omicron to antibody neutralization. Nature. 2022;602: 671–675. doi:10.1038/s41586-021-04389-z

9. Liu J, Wu Y, Gao GF. A Structural Voyage Toward the Landscape of Humoral and Cellular Immune Escapes of SARS-CoV-2. Immunological Reviews. 2025;330: e70000. doi:10.1111/imr.70000

10. van Dorp C, Goldberg E, Ke R, Hengartner N, Romero-Severson E. Global estimates of the fitness advantage of SARS-CoV-2 variant Omicron. Virus Evolution. 2022;8: veac089. doi:10.1093/ve/veac089

11. Puhach O, Meyer B, Eckerle I. SARS-CoV-2 viral load and shedding kinetics. Nat Rev Microbiol. 2023;21: 147–161. doi:10.1038/s41579-022-00822-w

12. Hui KPY, Ho JCW, Cheung M, Ng K, Ching RHH, Lai K, et al. SARS-CoV-2 Omicron variant replication in human bronchus and lung ex vivo. Nature. 2022;603: 715–720. doi:10.1038/s41586-022-04479-6

13. Barut GT, Halwe NJ, Taddeo A, Kelly JN, Schön J, Ebert N, et al. The spike gene is a major determinant for the SARS-CoV-2 Omicron-BA.1 phenotype. Nat Commun. 2022;13: 5929. doi:10.1038/s41467-022-33632-y

14. Shi G, Li T, Lai KK, Johnson RF, Yewdell JW, Compton AA. Omicron Spike confers enhanced infectivity and interferon resistance to SARS-CoV-2 in human nasal tissue | Nature Communications. Nature Communications. 2024;15: 889. doi:10.1038/s41467-024-45075-8

15. Tanneti NS, Patel AK, Tan LH, Marques AD, Perera RAPM, Sherrill-Mix S, et al. Comparison of SARS-CoV-2 variants of concern in primary human nasal cultures demonstrates Delta as most cytopathic and Omicron as fastest replicating. mBio. 2024;0: e03129–23. doi:10.1128/mbio.03129-23

16. Wolter N, Jassat W, Walaza S, Welch R, Moultrie H, Groome M, et al. Early assessment of the clinical severity of the SARS-CoV-2 omicron variant in South Africa: a data linkage study. The Lancet. 2022;399: 437–446. doi:10.1016/S0140-6736(22)00017-4

17. Pascall DJ, Vink E, Blacow R, Bulteel N, Campbell A, Campbell R, et al. Directions of change in intrinsic case severity across successive SARS-CoV-2 variant waves have been inconsistent. Journal of Infection. 2023;87: 128–135. doi:10.1016/j.jinf.2023.05.019

18. Armando F, Beythien G, Kaiser FK, Allnoch L, Heydemann L, Rosiak M, et al. SARS-CoV-2 Omicron variant causes mild pathology in the upper and lower respiratory tract of hamsters. Nat Commun. 2022;13: 3519. doi:10.1038/s41467-022-31200-y

19. de Melo GD, Perraud V, Alvarez F, Vieites-Prado A, Kim S, Kergoat L, et al. Neuroinvasion and anosmia are independent phenomena upon infection with SARS-CoV-2 and its variants. Nat Commun. 2023;14: 4485. doi:10.1038/s41467-023-40228-7

20. Rosenke K, Giffin A, Kaiser F, Altynova E, Mukesh R, Flagg M, et al. Pathogenesis of bovine H5N1 clade 2.3.4.4b infection in Macaques. Nature. 2025; 1–3. doi:10.1038/s41586-025-08609-8

21. Lamers MM, Mykytyn AZ, Breugem TI, Groen N, Knoops K, Schipper D, et al. SARS-CoV-2 Omicron efficiently infects human airway, but not alveolar epithelium. bioRxiv; 2022. p. 2022.01.19.476898. doi:10.1101/2022.01.19.476898

22. Masui A, Hashimoto R, Matsumura Y, Yamamoto T, Nagao M, Noda T, et al. Micro-patterned culture of iPSC-derived alveolar and airway cells distinguishes SARS-CoV-2 variants. Stem Cell Reports. 2024;19: 545–561. doi:10.1016/j.stemcr.2024.02.011

23. Hoffmann M, Krüger N, Schulz S, Cossmann A, Rocha C, Kempf A, et al. The Omicron variant is highly resistant against antibody-mediated neutralization: Implications for control of the COVID-19 pandemic. Cell. 2022;185: 447–456.e11. doi:10.1016/j.cell.2021.12.032

24. McCallum M, Czudnochowski N, Rosen LE, Zepeda SK, Bowen JE, Walls AC, et al. Structural basis of SARS-CoV-2 Omicron immune evasion and receptor engagement. Science. 2022;375: 864–868. doi:10.1126/science.abn8652

25. Parsons RJ, Acharya P. Evolution of the SARS-CoV-2 Omicron spike. Cell Reports. 2023;42: 113444. doi:10.1016/j.celrep.2023.113444

26. Wu C-T, Lidsky PV, Xiao Y, Cheng R, Lee IT, Nakayama T, et al. SARS-CoV-2 replication in airway epithelia requires motile cilia and microvillar reprogramming. Cell. 2023;186: 112–130.e20. doi:10.1016/j.cell.2022.11.030

27. Meng B, Abdullahi A, Ferreira IATM, Goonawardane N, Saito A, Kimura I, et al. Altered TMPRSS2 usage by SARS-CoV-2 Omicron impacts infectivity and fusogenicity. Nature. 2022;603: 706–714. doi:10.1038/s41586-022-04474-x

28. Willett BJ, Grove J, MacLean OA, Wilkie C, De Lorenzo G, Furnon W, et al. SARS-CoV-2 Omicron is an immune escape variant with an altered cell entry pathway. Nat Microbiol. 2022;7: 1161–1179. doi:10.1038/s41564-022-01143-7

29. Guo K, Barrett BS, Morrison JH, Mickens KL, Vladar EK, Hasenkrug KJ, et al. Interferon resistance of emerging SARS-CoV-2 variants. Proceedings of the National Academy of Sciences. 2022;119: e2203760119. doi:10.1073/pnas.2203760119

30. Nchioua R, Schundner A, Klute S, Koepke L, Hirschenberger M, Noettger S, et al. Reduced replication but increased interferon resistance of SARS-CoV-2 Omicron BA.1. Life Sci Alliance. 2023;6: e202201745. doi:10.26508/lsa.202201745

31. Reuschl A-K, Thorne LG, Whelan MVX, Ragazzini R, Furnon W, Cowton VM, et al. Evolution of enhanced innate immune suppression by SARS-CoV-2 Omicron subvariants. Nat Microbiol. 2024; 1–13. doi:10.1038/s41564-023-01588-4

32. Robinot R, Hubert M, de Melo GD, Lazarini F, Bruel T, Smith N, et al. SARS-CoV-2 infection induces the dedifferentiation of multiciliated cells and impairs mucociliary clearance. Nat Commun. 2021;12: 4354. doi:10.1038/s41467-021-24521-x

33. Elad D, Wolf M, Keck T. Air-conditioning in the human nasal cavity. Respiratory Physiology & Neurobiology. 2008;163: 121–127. doi:10.1016/j.resp.2008.05.002

34. Keck T, Leiacker R, Riechelmann H, Rettinger G. Temperature Profile in the Nasal Cavity. The Laryngoscope. 2000;110: 651–654. doi:10.1097/00005537-200004000-00021

35. Rajah MM, Hubert M, Bishop E, Saunders N, Robinot R, Grzelak L, et al. SARS-CoV-2 Alpha, Beta, and Delta variants display enhanced Spike-mediated syncytia formation. The EMBO Journal. 2021;40: e108944. doi:10.15252/embj.2021108944

36. Ziegler CGK, Miao VN, Owings AH, Navia AW, Tang Y, Bromley JD, et al. Impaired local intrinsic immunity to SARS-CoV-2 infection in severe COVID-19. Cell. 2021;184: 4713–4733.e22. doi:10.1016/j.cell.2021.07.023

37. Rodero MP, Decalf J, Bondet V, Hunt D, Rice GI, Werneke S, et al. Detection of interferon alpha protein reveals differential levels and cellular sources in disease. Journal of Experimental Medicine. 2017;214: 1547–1555. doi:10.1084/jem.20161451

38. Sposito B, Broggi A, Pandolfi L, Crotta S, Clementi N, Ferrarese R, et al. The interferon landscape along the respiratory tract impacts the severity of COVID-19. Cell. 2021;184: 4953–4968.e16. doi:10.1016/j.cell.2021.08.016

39. Stanifer ML, Pervolaraki K, Boulant S. Differential Regulation of Type I and Type III Interferon Signaling. International Journal of Molecular Sciences. 2019;20: 1445. doi:10.3390/ijms20061445

40. Zhang Q, Bastard P, Cobat A, Casanova J-L. Human genetic and immunological determinants of critical COVID-19 pneumonia. Nature. 2022;603: 587–598. doi:10.1038/s41586-022-04447-0

41. Bastard P, Gervais A, Le Voyer T, Rosain J, Philippot Q, Manry J, et al. Autoantibodies neutralizing type I IFNs are present in ∼4% of uninfected individuals over 70 years old and account for ∼20% of COVID-19 deaths. Science Immunology. 2021;6: eabl4340. doi:10.1126/sciimmunol.abl4340

42. Caobi A, Saeed M. Upping the ante: enhanced expression of interferon-antagonizing ORF6 and ORF9b proteins by SARS-CoV-2 variants of concern. Current Opinion in Microbiology. 2024;79: 102454. doi:10.1016/j.mib.2024.102454

43. Le Pen J, Rice CM. The antiviral state of the cell: lessons from SARS-CoV-2. Current Opinion in Immunology. 2024;87: 102426. doi:10.1016/j.coi.2024.102426

44. Otter CJ, Renner DM, Fausto A, Tan LH, Cohen NA, Weiss SR. Interferon signaling in the nasal epithelium distinguishes among lethal and common cold coronaviruses and mediates viral clearance. Proceedings of the National Academy of Sciences. 2024;121: e2402540121. doi:10.1073/pnas.2402540121

45. Thorne LG, Bouhaddou M, Reuschl A-K, Zuliani-Alvarez L, Polacco B, Pelin A, et al. Evolution of enhanced innate immune evasion by SARS-CoV-2. Nature. 2022;602: 487–495. doi:10.1038/s41586-021-04352-y

46. Mears HV, Young GR, Sanderson T, Harvey R, Barrett-Rodger J, Penn R, et al. Emergence of SARS-CoV-2 subgenomic RNAs that enhance viral fitness and immune evasion. PLOS Biology. 2025;23: e3002982. doi:10.1371/journal.pbio.3002982

47. Laine L, Skön M, Väisänen E, Julkunen I, Österlund P. SARS-CoV-2 variants Alpha, Beta, Delta and Omicron show a slower host cell interferon response compared to an early pandemic variant. Front Immunol. 2022;13. doi:10.3389/fimmu.2022.1016108

48. Bouhaddou M, Reuschl A-K, Polacco BJ, Thorne LG, Ummadi MR, Ye C, et al. SARS-CoV-2 variants evolve convergent strategies to remodel the host response. Cell. 2023; S0092-8674(23)00915–7. doi:10.1016/j.cell.2023.08.026

49. Thorne LG, Reuschl A, Zuliani-Alvarez L, Whelan MVX, Turner J, Noursadeghi M, et al. SARS-CoV-2 sensing by RIG-I and MDA5 links epithelial infection to macrophage inflammation. The EMBO Journal. 2021;40: e107826. doi:10.15252/embj.2021107826

50. Decalf J, Desdouits M, Rodrigues V, Gobert F-X, Gentili M, Marques-Ladeira S, et al. Sensing of HIV-1 Entry Triggers a Type I Interferon Response in Human Primary Macrophages. Journal of Virology. 2017;91: 10.1128/jvi.00147-17. doi:10.1128/jvi.00147-17

51. Walsh JML, Miao VN, Owings AH, Tang Y, Bromley JD, Kazer SW, et al. Variants and vaccines impact nasal immunity over three waves of SARS-CoV-2. Nat Immunol. 2025; 1–14. doi:10.1038/s41590-024-02052-z

52. Resnick JD, Beer MA, Pekosz A. Early Transcriptional Responses of Human Nasal Epithelial Cells to Infection with Influenza A and SARS-CoV-2 Virus Differ and Are Influenced by Physiological Temperature. Pathogens. 2023;12: 480. doi:10.3390/pathogens12030480

53. V’kovski P, Gultom M, Kelly JN, Steiner S, Russeil J, Mangeat B, et al. Disparate temperature-dependent virus-host dynamics for SARS-CoV-2 and SARS-CoV in the human respiratory epithelium. PLoS Biol. 2021;19: e3001158. doi:10.1371/journal.pbio.3001158

54. Laporte M, Raeymaekers V, Van Berwaer R, Vandeput J, Marchand-Casas I, Thibaut H-J, et al. The SARS-CoV-2 and other human coronavirus spike proteins are fine-tuned towards temperature and proteases of the human airways. PLoS Pathog. 2021;17: e1009500. doi:10.1371/journal.ppat.1009500

55. Otter CJ, Fausto A, Tan LH, Khosla AS, Cohen NA, Weiss SR. Infection of primary nasal epithelial cells differentiates among lethal and seasonal human coronaviruses. Proceedings of the National Academy of Sciences. 2023;120: e2218083120. doi:10.1073/pnas.2218083120

56. Heron I, Berg K. The actions of interferon are potentiated at elevated temperature. Nature. 1978;274: 508–510. doi:10.1038/274508a0

57. Lane WC, Dunn MD, Gardner CL, Lam LKM, Watson AM, Hartman AL, et al. The Efficacy of the Interferon Alpha/Beta Response versus Arboviruses Is Temperature Dependent. mBio. 2018;9: 10.1128/mbio.00535-18. doi:10.1128/mbio.00535-18

58. Wang Q, Anang S, Iketani S, Guo Y, Liu L, Katsamba PS, et al. Functional properties of the spike glycoprotein of the emerging SARS-CoV-2 variant B.1.1.529. Cell Reports. 2022;39: 110924. doi:10.1016/j.celrep.2022.110924

59. Gobeil SM-C, Henderson R, Stalls V, Janowska K, Huang X, May A, et al. Structural diversity of the SARS-CoV-2 Omicron spike. Molecular Cell. 2022;82: 2050–2068.e6. doi:10.1016/j.molcel.2022.03.028

60. Dufloo J, Sanjuán R. Temperature impacts SARS-CoV-2 spike fusogenicity and evolution. mBio. 2024;15: e03360–23. doi:10.1128/mbio.03360-23

61. Bisht K, te Velthuis AJW. Decoding the Role of Temperature in RNA Virus Infections. mBio. 2022;13: e02021–22. doi:10.1128/mbio.02021-22

62. Chan JF-W, Huang X, Hu B, Chai Y, Shi H, Zhu T, et al. Altered host protease determinants for SARS-CoV-2 Omicron. Science Advances. 2023;9: eadd3867. doi:10.1126/sciadv.add3867

63. Mykytyn AZ, Breugem TI, Geurts MH, Beumer J, Schipper D, van Acker R, et al. SARS-CoV-2 Omicron entry is type II transmembrane serine protease-mediated in human airway and intestinal organoid models. Journal of Virology. 2023;0: e00851–23. doi:10.1128/jvi.00851-23

64. Becker ME, Martin-Sancho L, Simons LM, McRaven MD, Chanda SK, Hultquist JF, et al. Live imaging of airway epithelium reveals that mucociliary clearance modulates SARS-CoV-2 spread. Nat Commun. 2024;15: 9480. doi:10.1038/s41467-024-53791-4

65. Li Q, Vijaykumar K, Phillips SE, Hussain SS, Huynh NV, Fernandez-Petty CM, et al. Mucociliary transport deficiency and disease progression in Syrian hamsters with SARS-CoV-2 infection. JCI Insight. 2023;8: e163962. doi:10.1172/jci.insight.163962

66. Vijaykumar K, Leung HM, Barrios A, Fernandez-Petty CM, Solomon GM, Hathorne HY, et al. COVID-19 Causes Ciliary Dysfunction as Demonstrated by Human Intranasal Micro-Optical Coherence Tomography Imaging. Am J Respir Cell Mol Biol. 2023;69: 592–595. doi:10.1165/rcmb.2023-0177LE

67. Bustamante-Marin XM, Ostrowski LE. Cilia and Mucociliary Clearance. Cold Spring Harb Perspect Biol. 2017;9: a028241. doi:10.1101/cshperspect.a028241

68. Cheng L, Li Z, Wu H, Li F, Qiu Y, Wang T, et al. Clinical and pathogen features of COVID-19-associated infections during an Omicron strain outbreak in Guangzhou, China. Microbiology Spectrum. 2024;12: e03406–23. doi:10.1128/spectrum.03406-23

69. Murakami Y, Nozaki Y, Morosawa M, Toyama M, Ogashiwa H, Ueda T, et al. Difference in the impact of coinfections and secondary infections on antibiotic use in patients hospitalized with COVID-19 between the Omicron-dominant period and the pre-Omicron period. Journal of Infection and Chemotherapy. 2024;30: 853–859. doi:10.1016/j.jiac.2024.02.026

70. Planchais C, Fernández I, Bruel T, de Melo GD, Prot M, Beretta M, et al. Potent human broadly SARS-CoV-2–neutralizing IgA and IgG antibodies effective against Omicron BA.1 and BA.2. Journal of Experimental Medicine. 2022;219: e20220638. doi:10.1084/jem.20220638

71. Llibre A, Bilek N, Bondet V, Darboe F, Mbandi SK, Penn-Nicholson A, et al. Plasma Type I IFN Protein Concentrations in Human Tuberculosis. Front Cell Infect Microbiol. 2019;9. doi:10.3389/fcimb.2019.00296

72. Andrews S. FastQC A Quality Control tool for High Throughput Sequence Data. In: Babraham Bioinformatics [Internet]. 2010 [cited 9 Jan 2025]. Available: https://www.bioinformatics.babraham.ac.uk/projects/fastqc/

73. Chen S, Zhou Y, Chen Y, Gu J. fastp: an ultra-fast all-in-one FASTQ preprocessor. Bioinformatics. 2018;34: i884–i890. doi:10.1093/bioinformatics/bty560

74. Kim D, Paggi JM, Park C, Bennett C, Salzberg SL. Graph-based genome alignment and genotyping with HISAT2 and HISAT-genotype. Nat Biotechnol. 2019;37: 907–915. doi:10.1038/s41587-019-0201-4

75. Tarasov A, Vilella AJ, Cuppen E, Nijman IJ, Prins P. Sambamba: fast processing of NGS alignment formats. Bioinformatics. 2015;31: 2032–2034. doi:10.1093/bioinformatics/btv098

76. Liao Y, Smyth GK, Shi W. featureCounts: an efficient general purpose program for assigning sequence reads to genomic features. Bioinformatics. 2014;30: 923–930. doi:10.1093/bioinformatics/btt656

77. Wu T, Hu E, Xu S, Chen M, Guo P, Dai Z, et al. clusterProfiler 4.0: A universal enrichment tool for interpreting omics data. The Innovation. 2021;2: 100141. doi:10.1016/j.xinn.2021.100141

78. Chen EY, Tan CM, Kou Y, Duan Q, Wang Z, Meirelles GV, et al. Enrichr: interactive and collaborative HTML5 gene list enrichment analysis tool. BMC Bioinformatics. 2013;14: 128. doi:10.1186/1471-2105-14-128

79. Wickham H. ggplot2: Elegant Graphics for Data Analysis (3e). Springer; 2016. Available: https://ggplot2-book.org/

