## Supplemental Figures S1 to S5 and Sup. Table 1 for "Stealth replication of SARS-CoV-2 Omicron in the nasal epithelium at physiological temperature"

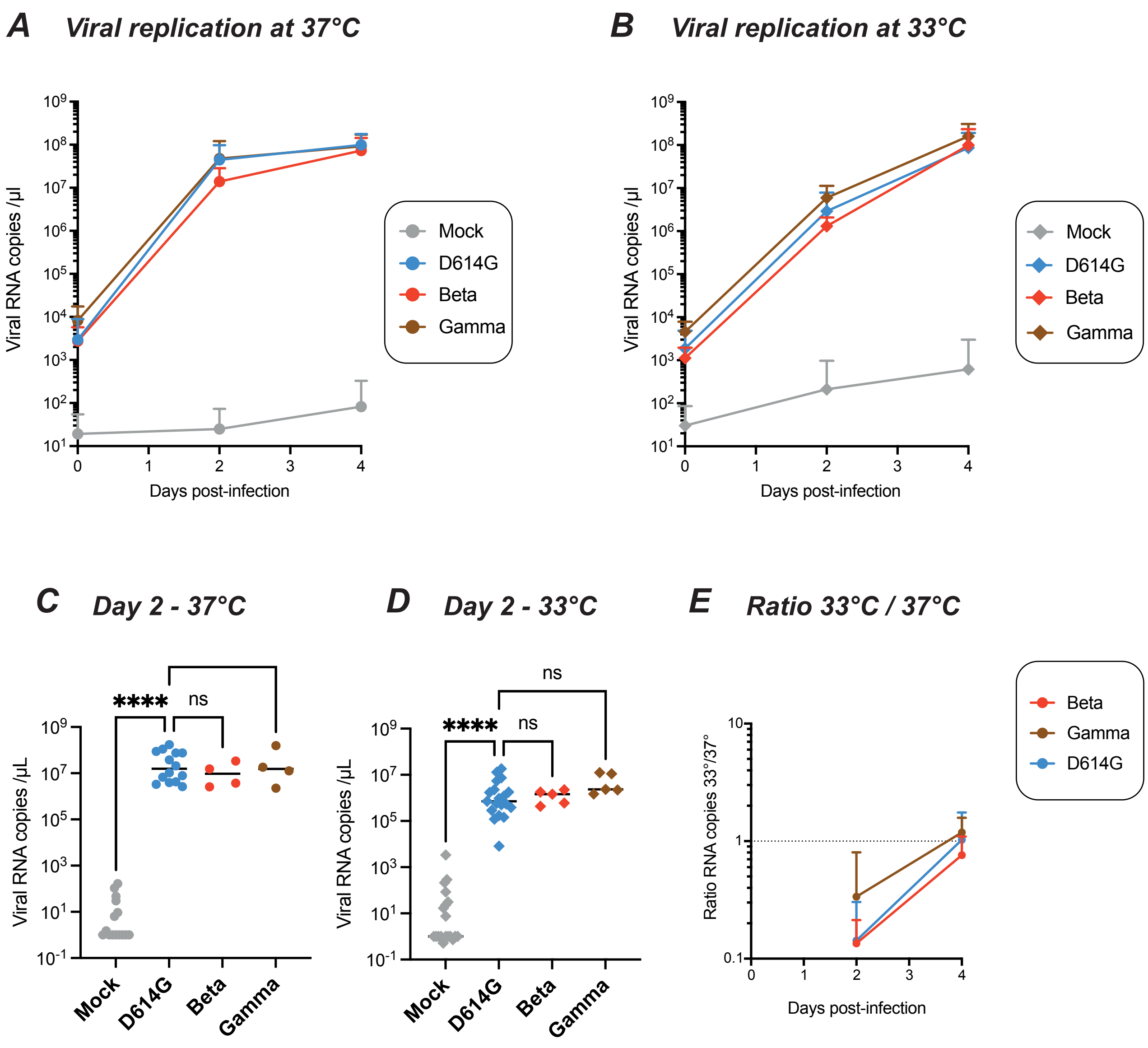


***Figure S1: Temperature-dependent replication of the Beta and Gamma SARS-CoV-2 variants in reconstructed human nasal epithelia.***

*The viral load in apical supernatants was quantified by RT-qPCR for samples infected by the following SARS-CoV-2 lineages: D614G, Beta, and Gamma. (A, B) Kinetics of viral replication at 37°C (A) and at 33°C (B), with means and SD reported (n=4 to 14 independent samples per point). (C,D) Comparison of apical viral loads measured at 2 days post infection (dpi) at 37°C (C) and 33°C (D). Medians are reported. (E) The ratio of viral RNA copies measured at 33°C to those measured at 37°C is shown at different dpi, with means and SD reported (n=4 to 14 independent samples per point).(C, D, E) Statistical comparisons were done between the D614G reference and all other variants or the Mock condition, using the Kruskal-Wallis test with Dunn's correction; ns: not significant; **** P<0.0001.*

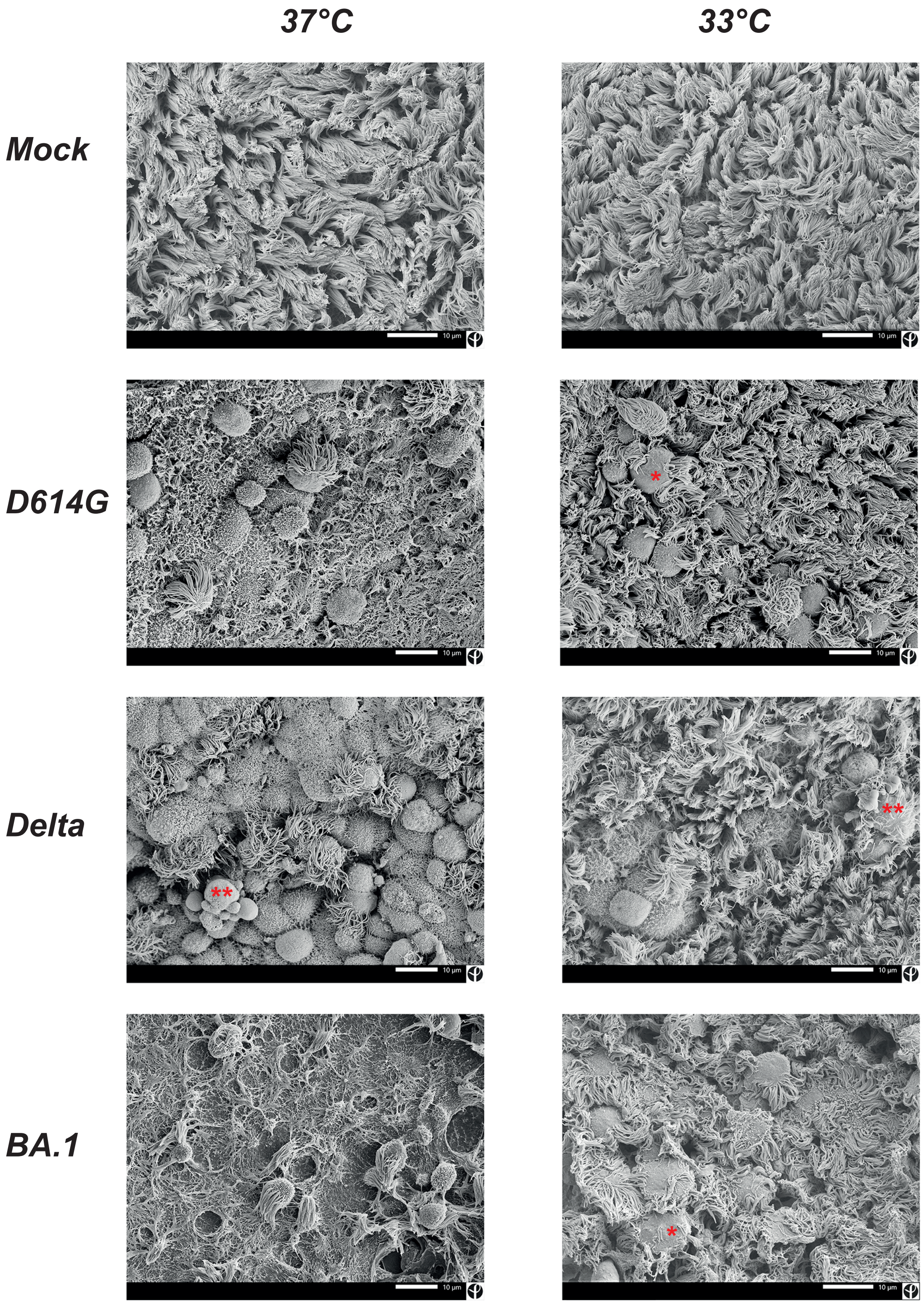


***Figure S2: Infection by SARS-CoV-2 variants induce a loss of motile cilia.***
*Scanning electron microscopy images of reconstructed nasal epithelia at 4 days post-infection. The epithelia were infected at 37°C (left) or at 33°C (right) with the D614G, Delta or Omicron, BA.1 variant (shown in rows 2, 3, 4, respectively) or were mock-infected (first row). The scale bar represents 10 µm.*
*Red symbols: * membrane protrusion surrounding cilia; ** blebbing cell.*

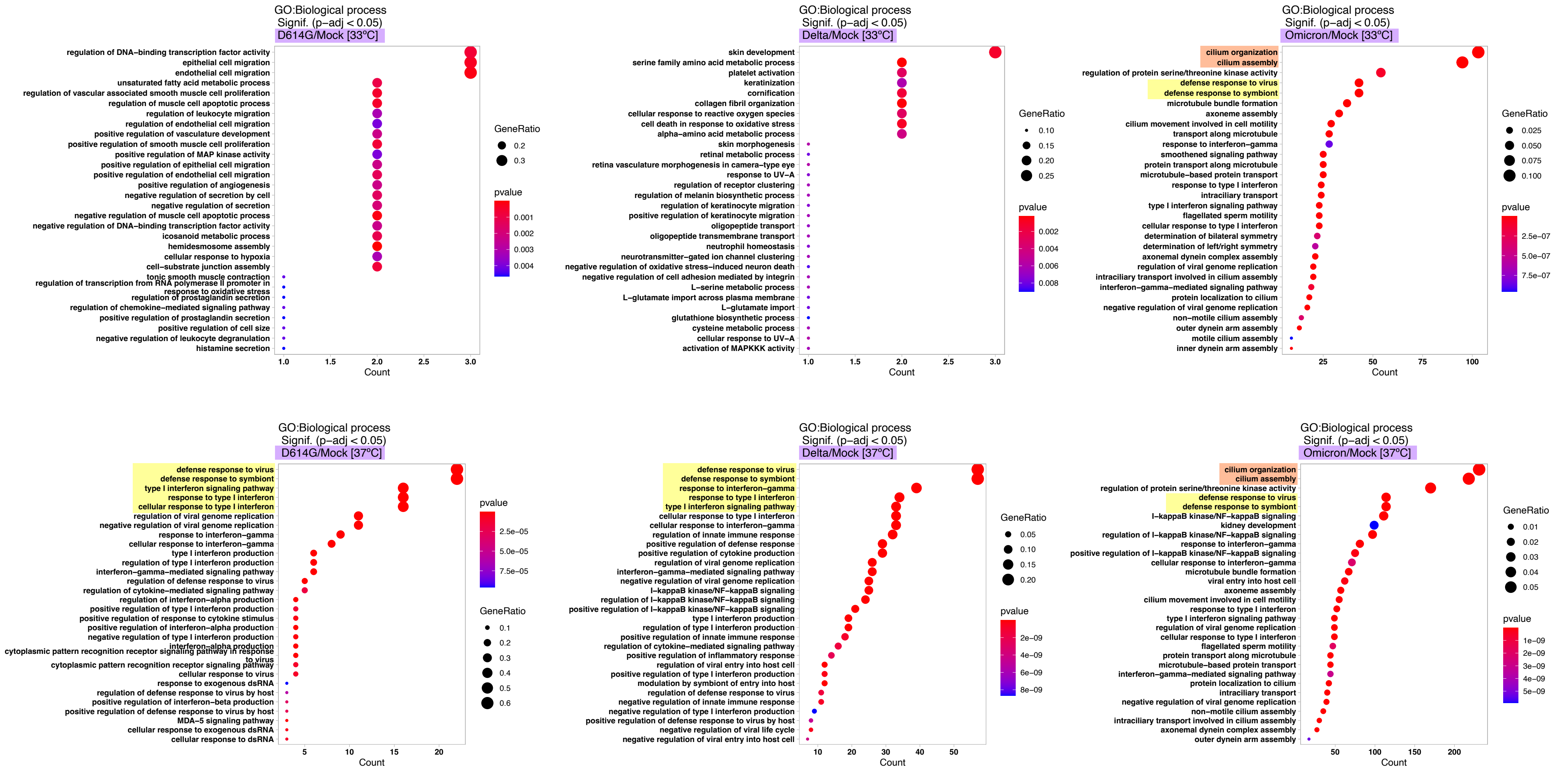


***Figure S3: Functional enrichment analysis of transcriptomes from reconstructed nasal epithelia at day 2 post-infection.*** *The significantly dysregulated gene ontology (GO) terms are reported for epithelial samples infected at 33°C (top) or 37°C (bottom) with the variants D614G (left), delta (middle), and Omicron BA.1 (right). The number of differentially expressed genes (DEGs) belonging to each GO term is reported on the x axis. The size and color of symbols correspond to the gene ratio and p value for each GO term, respectively. Among the 5 top dysregulated GO terms, those corresponding to "cilium organization" and "defense response" are highlighted in orange and yellow, respectively.*

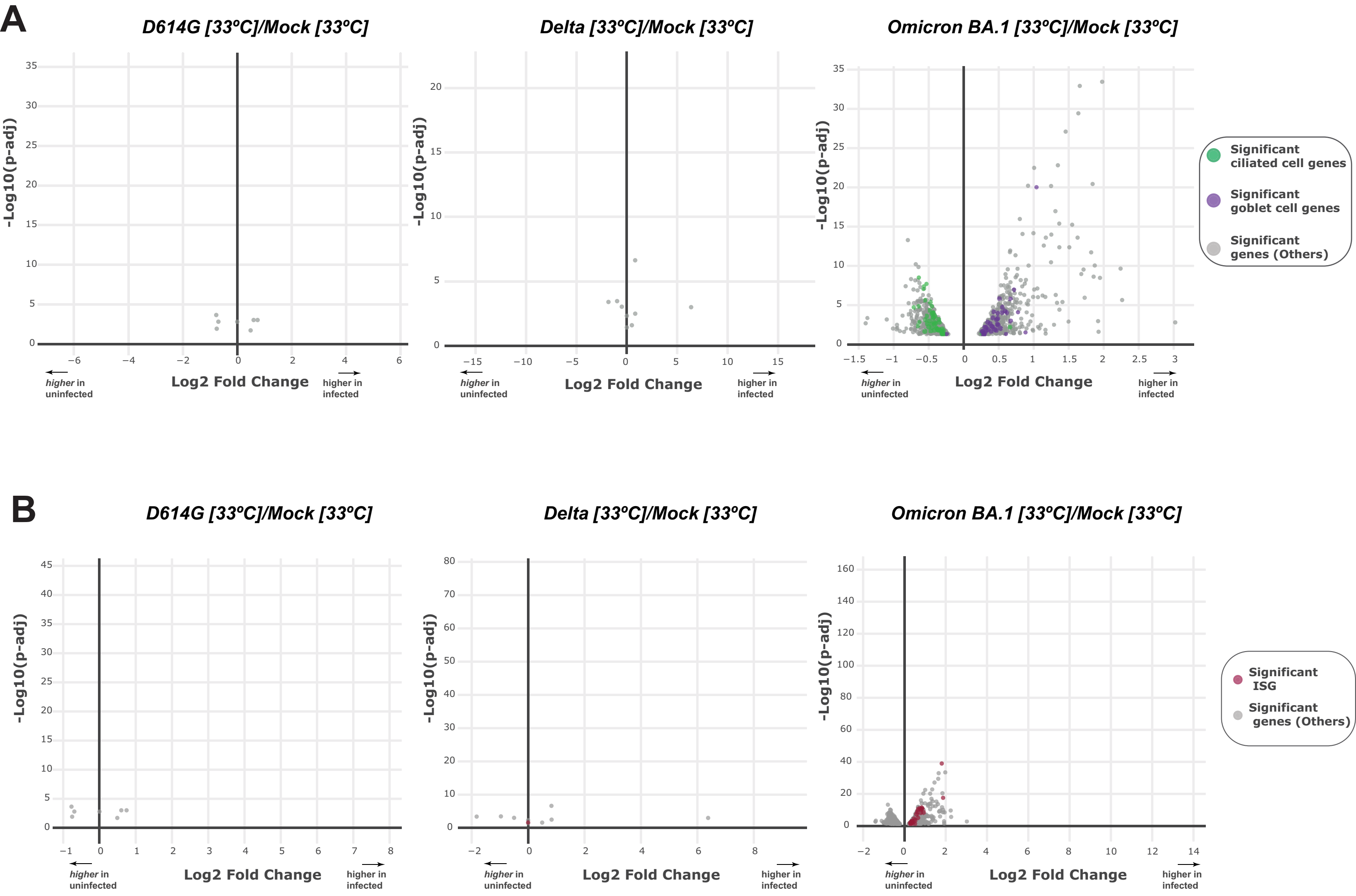


***Figure S4: Limited changes in transcriptional profiles of reconstructed nasal epithelia infected at 33°C.***
*(A) Volcano-plots depicting the differential regulation of ciliated cell genes (green) and goblet cell genes (purple) at 33°C in variant-infected samples compared to mock-infected samples at 2 dpi. The log2 fold change in gene expression is shown on the x axis and the adjusted p value on the y axis. Genes overexpressed in infected samples are distributed to the right on the x axis. Significant DEGs that do not belong to the ciliated cell or goblet cell category are represented in grey. (B) Volcano-plots depicting the differential regulation of ISGs (red) at 33°C in variant-infected samples compared to mock-infected samples at 2 dpi. The representation is similar to that in (A).*

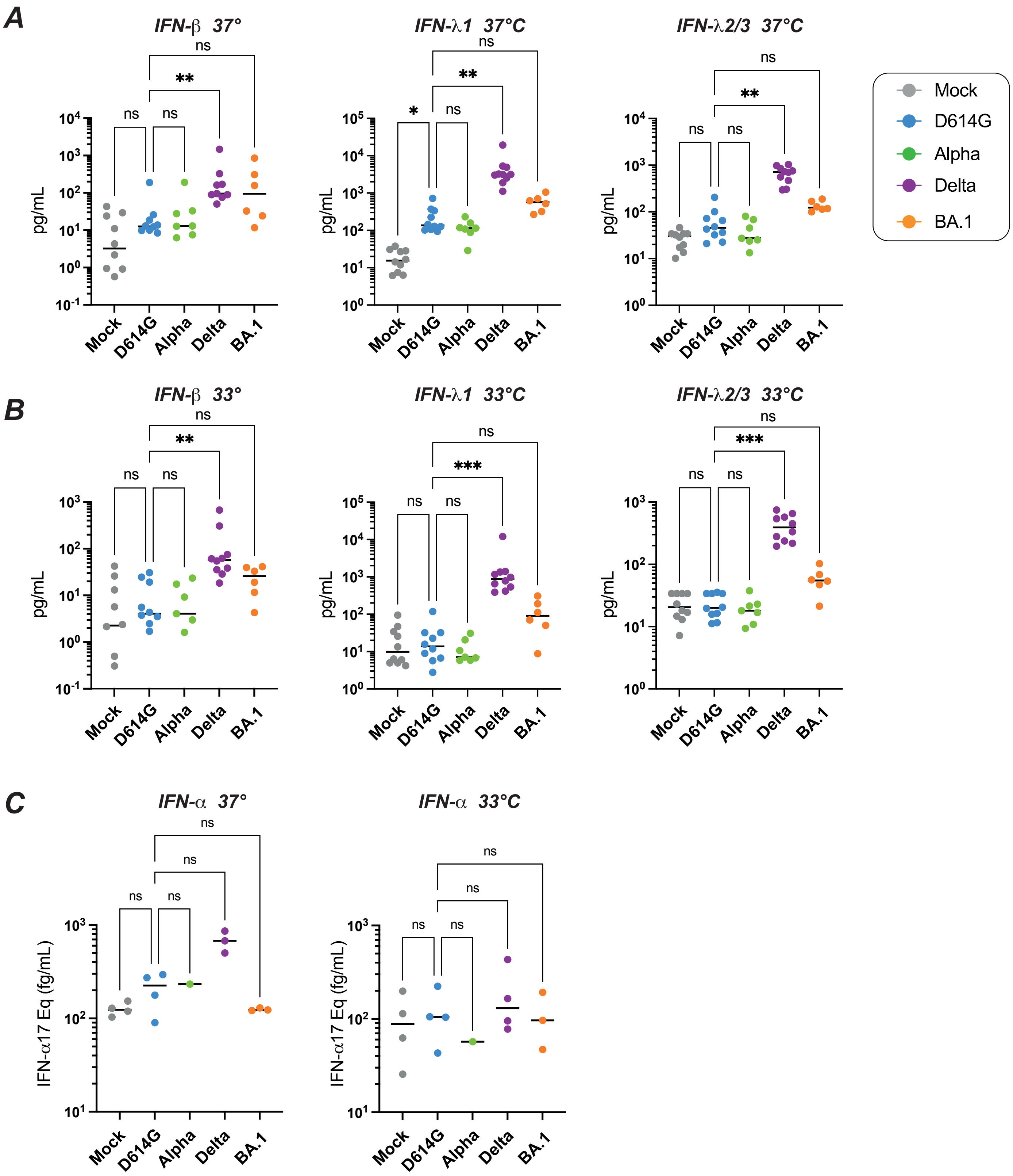


***Figure S5: Induction of type I and type III IFNs by SARS-CoV-2 variants at day 4 post-infection.***

*(A) The induction of IFN-β (left), IFN-λ1 (middle) and IFN-λ2/3 (right) release in apical supernatants of reconstructed nasal epithelia infected at 37°C was measured by digital ELISA (IFN-β) and Legendplex assays (IFN-λ). (B) Similar graphs as in (A), but for infections carried out at 33°C. (C) The induction of IFN-α release in apical supernatants of reconstructed nasal epithelia infected at 37°C (left) and 33°C (right) was measured by digital ELISA, with results expressed as IFN-α17 equivalent. (A-C) Medians are reported. Statistical comparisons were done between the D614G reference variants and all the other variants tested, using the Kruskal-Wallis test with Dunn's correction; ns: not significant; ns: not significant; * P<0.05; ** P<0.01; *** P<0.001.*

**Supplementary Table 1: Viral Stocks used in the study**

| Variant of concern | Strain derivation | Gisaid/EVAg | Reference | DOI |
|---|---|---|---|---|
| **Wuhan** | BetaCoV/France/IDF00372/2020 | Ref-SKU: 014V-0389[illegible] | Robinot R et al, Nat Comm. 2021 | doi: 10.1038/s41467-021-24521-x |
| **D614G** | hCoV-19/France/GE1973/2020* | EPI_ISL_414631 | Planas D et al, Nat Med 2021 | doi: 10.1038/s41591-021-01318-5 |
| **Alpha (B.1.1.7)** | hCoV-19/France/CVL-SC719/2020 | EPI_ISL_735391 | Planas D et al, Nat Med 2021 | doi: 10.1038/s41591-021-01318-5 |
| **Beta (B.1.351)** | hCoV-19/France/IDF-IPP00078/2021 | EPI_ISL_964916 | Planas D et al, Nat Med 2021 | doi: 10.1038/s41591-021-01318-5 |
| **Gamma (P.1)** | hCoV-19/Japan/TY7-501/2021 | EPI_ISL_833366 | Betton M. et al., Clin Inf Dis 2021 | doi: 10.1093/cid/ciab308 |
| **Delta (B.1.617)** | hCoV-19/France/IDF-APHP-HEGP-20-23-2131905084/2021 | EPI_ISL_2029113 | Planas D et al, Nature 2021 | doi: 10.1038/s41586-021-03777-9 |
| **Omicron (BA.1)** | hCoV-19/Belgium/rega-20174/2021 | EPI_ISL_6794907 | Planas D et al, Nature 2022 | doi: 10.1038/s41586-021-04389-z |

Notes *Viral isolate supplied by the National Reference Centre for Respiratory Viruses hosted by Institut Pasteur
